# Hydrogen-bonded organic framework nanotransducers enabled sono-optogenetics for Parkinsonian rats

**DOI:** 10.1101/2025.01.03.631271

**Authors:** Wenliang Wang, Ilya Pyatnitskiy, Yanshu Shi, Kai Wing Kevin Tang, Yi Xie, Thomas Wynn, Xiangping Liu, Ju-Chun Hsieh, Jinmo Jeong, Weilong He, Brinkley Artman, Anakaren Romero Lozano, Xi Shi, Arjun Sangani, Lief Fenno, Samantha Santacruz, Banglin Chen, Huiliang Wang

## Abstract

Cell-type-specific activation of parvalbumin (PV)-expressing neurons in the external globus pallidus (GPe) through optogenetics has shown promise in facilitating long-lasting movement dysfunction recovery in mice with Parkinson’s disease. However, its translational potential is hindered by adverse effects stemming from the invasive implantation of optical fibers into the brain. In this study, we have developed a non-invasive optogenetics approach, utilizing focused ultrasound-triggered mechanoluminescent nanotransducers to enable remote photon delivery deep in the brain for genetically targeted neuromodulation. These mechanoluminescent nanotransducers consist of sonosensitized hydrogen-bonded frameworks and chemiluminescent L012, serving as a nanoscale light source through ultrasound-induced cascade reactions. This system offers high ultrasound-triggered brightness and long-lasting light emission, facilitating repeatable deep brain stimulation. Our sono-optogenetics technology demonstrated effective modulation in the mouse motor cortex for limb motion control and activation of PV-GPe neurons for rescuing movement dysfunction over time in dopamine-depleted Parkinson’s disease rats. This approach demonstrates the pathway for achieving genetically targeted and non-invasive neuromodulation for long-lasting treatment of Parkinson’s disease, towards non-human primate models and clinical applications.

---

Parkinson’s disease (PD) results in significant motor dysfunctions, tremor, rigidity, and cognitive impairments in the later stages, presenting a crucial challenge to the global healthcare community.^1,2^ Deep brain stimulation (DBS) has emerged as an effective clinical intervention for transiently alleviating motor symptoms in PD patients, through high-frequency electrical stimulation of the subthalamic nucleus or globus pallidus interna with implanted microelectrodes.^3,4^ However, the high cost and significant surgical risks, including hemorrhage and infection, typically limit the extensive application of DBS in clinical PD treatment.^5^ In addition, a significant limitation of the existing DBS therapies is that they treat the symptoms of the disease, but do not correct the underlying circuit dysfunction responsible for the symptoms.^6–8^ As a result, the long-term alleviation of these symptoms is limited, and the disease prognosis often remains unchanged despite the invasive nature of DBS. Furthermore, various circuits are recruited during electrical stimulation for PD treatment, leading to non-specific stimulations and potential widespread prodromic and antidromic changes in brain activity.^9,10^ Consequently, there is a pressing demand for non-invasive neural circuit-selective stimulation technologies for effective PD treatment.

Optogenetics provides spatiotemporal and cell-type specific control of neural circuits that have opened the door for precisely targeted PD treatment.^9,11–14^ Recent advancements have demonstrated that optogenetic activation of parvalbumin-expressing globus pallidus neurons (PV-GPe) can effectively restore movement and reverse pathological activity in dopamine-deficient mice over time, providing a precise circuit targeting, long-lasting PD treatment methods with minimum changes in brain activity.^8^ While significant strides have been made in Parkinson’s disease management through cell-type-specific optogenetics, its clinical application remains still constrained by surgery-related adverse effects reminiscent of those observed in DBS, stemming from invasive fiber implantations.^3,9,13,15^ The permanent implantation of fiber or electrodes could cause inconvenience to daily life and increase the risk of postoperative infection. Therefore, non-invasive and implantation-free optogenetics holds great potential for clinical treatments of PD.

Focused ultrasound (FUS)-triggered mechanoluminescent nanoparticles offer a promising avenue for non-invasive light delivery in deep brain regions, leveraging ultrasound’s exceptional tissue penetration and established clinical safety.^16–18^ Inorganic nanoparticles were first demonstrated to generate light upon FUS stimulation, and applied for the non-invasive mouse motor cortex modulation, named sono-optogenetics.^16,19,20^ Yet, their application has been encumbered by the need for prerequisite charging and potential biocompatibility issues with inorganic nanoparticles.^21–24^ More recently, the emergence of organic nanoparticles, driven by ultrasound-induced cascade reactions between sonosensitizers and chemiluminescent reagents in liposomes, has enabled the achievement of active light emission without charging.^18,25^ This innovation holds promise for sustainable and repeatable sono-optogenetics, bolstered by improved biocompatibility and system simplicity.^18^ Nonetheless, the restricted drug loading capacity within these nanoparticles has curtailed the efficiency of cascade reactions, resulting in diminished luminescence intensity and duration under ultrasound stimulation.^17,18,26^ The development of mechanoluminescent nanoparticles with higher photon emission and more repeatable stimulation remains a primary challenge in achieving effective deep brain sono-optogenetics for disease treatments.

Hydrogen-bonded organic frameworks (HOFs), assembled through intermolecular multivalent hydrogen bonds and π-π stacking interactions of organic building units, have recently emerged as a promising class of porous materials with high structural homogeneity and programmability.^27,28^ They exhibit high drug loading capacity, biocompatibility, and functional programmability achieved by adjusting building units for various bioapplications.^29–31^ With this regard, we hypothesize that a HOF constructed from sonosensitizer building units could serve as porous carriers to load a high percentage of chemiluminescent reagents, acting as ultrasound-induced cascade reaction containers to achieve higher photon yields for deep brain sono-optogenetics. In addition, despite substantial advancements in ultrasound-activated mechanoluminescence materials recently, their application in the treatment of neurological disorders remains largely unexplored. As such, the integration of sono-optogenetics with the targeted modulation of PV-GPe neural circuits in a Parkinson’s disease rat model could herald new avenues for clinical therapeutic interventions for Parkinson’s disease.

In this study, we present the design of a sonosensitized HOF incorporating chemiluminescent L012, serving as a nanoscale light source to enable high intensity photon delivery over time with high temporal resolution. Furthermore, we demonstrated the targeted activation of PV-GPe neurons through these nanotransducers under FUS stimulation, ultimately ameliorating motor dysfunction in a Parkinson’s disease rat model. These findings establish the feasibility of non-invasive, genetically targeted deep brain neuromodulation technology through sono-optogenetics and underscore their potential as therapeutic techniques for the treatment of PD. We anticipate that sono-optogenetics will continue to evolve and find expanded applications in the treatment of various neurological diseases, offering a precise and biocompatible neuromodulation approach for clinical applications.

## Results and Discussion

### Development of sonosensitized HOF nanoparticles

A liposomal light source, based on the ultrasound-triggered cascade reaction between sonosensitizers and chemoluminescent L012, has been developed to achieved wireless photon delivery in the brain.^18^ However, the photon yield efficiency and long-lasting emission were limited by the restricted loading capacity of liposomal system. Porous HOF exhibited excellent biocompatibility and biosafety attributing to the metal-free nature compared with metal organic frameworks, making them a desirable candidate for drug delivery with excellent loading capacity.^30,32^ In addition, we could programmably design the organic building units of HOFs to create a well-defined sonosensitized delivery platform. We anticipate that the porous and stable HOF with desirable structures could serve as an efficient container for high-efficiency L012 loading and as a sonosensitizer for ROS generation, thereby enabling the activation of L012 for light emission under ultrasound stimulation.

Here, the 1,3,6,8-tetrakis(p-benzoic acid) pyrene (H4TBAPy) ligand, a planar molecule with a large aromatic fused ring and four carboxylate acid groups is designed to build the sonosensitized HOF (Sono-HOF, **Fig. 1a**). The narrow bandgap of the H4TBAPy ligand (1.723 eV) in its crystal form, as calculated using first principles density functional theory (DFT) in the Dmol3 code, is advantageous for promoting high-yield ROS generation under low-intensity ultrasound parameters compared to IR-780 (1.883 eV).^33^ This is because the narrow bandgap of the sonosensitizer dominates the minimal ultrasound energy required to generate electron-hole (e^-^-h^+^) pairs for ROS production.^34,35^ Additionally, the presence of four carboxylate acid groups conjugated with the fused ring contributes to the improved stability of the HOF in solution for long-term applications due to the high cohesive energy of frameworks.^36^ Then, nano-sized Sono-HOF particles were prepared through a solvent-exchange approach using H4TBAPy monomers.^30,36^ Powder X-ray diffraction confirmed the high crystallinity and phase purity of these Sono-HOF nanoparticles (**Supplementary Fig. 1a**). Each H4TBAPy unit formed interactions with four neighboring units through eight O-H⋯O hydrogen bonds, resulting in the formation of a two-dimensional layer.^30,37^ These layers then interacted with adjacent layers through π-π stacking interactions, self-assembling into three-dimensional frameworks.^30,37^ Experimental nitrogen adsorption tests at 77 K revealed that the Sono-HOF nanoparticles possessed a Brunauer−Emmett−Teller (BET) surface area of 497.5 m^2^ g^-1^ and a pore volume of 34.7 cm^3^ mol^-1^ (**Supplementary Fig. 1**). Transmission electron microscopy (TEM) showed regular multi-layered frameworks in these nanoparticles (**Fig. 1b**). Dynamic light scattering tests confirmed a well-distributed particle size with an average of 533.7 ± 14.9 nm (**Fig. 1c** and **Supplementary Table 1**). Of note, both the strong π-π stacking interactions from the extensive π-conjugated system and multivalent hydrogen bonding ensure the stability of the framework in solution. Remarkably, no significant changes in size or morphology were observed after 7 days of incubation, even under ultrasound stimulation (**Supplementary Fig. 2**). Additionally, the negative surface potential of these nanoparticles ensures their stability even in the presence of, as demonstrated in **Fig. 1c** and **Supplementary Table 1**. These sono-HOF nanoparticles exhibited excellent drug loading capacity, approximately ∼25 wt%, attributed to their high porosity, with no noticeable size changes observed after drug loading (**Supplementary Table 1**). To assess drug stability within the HOF nanoparticles, dye release tests were conducted. The results revealed that no dye was released under ultrasound stimulation and less than 10 % cargoes were released even after 36 h incubation, indicating long-term drug stability (**Supplementary Fig. 3**).

**Fig. 1.**
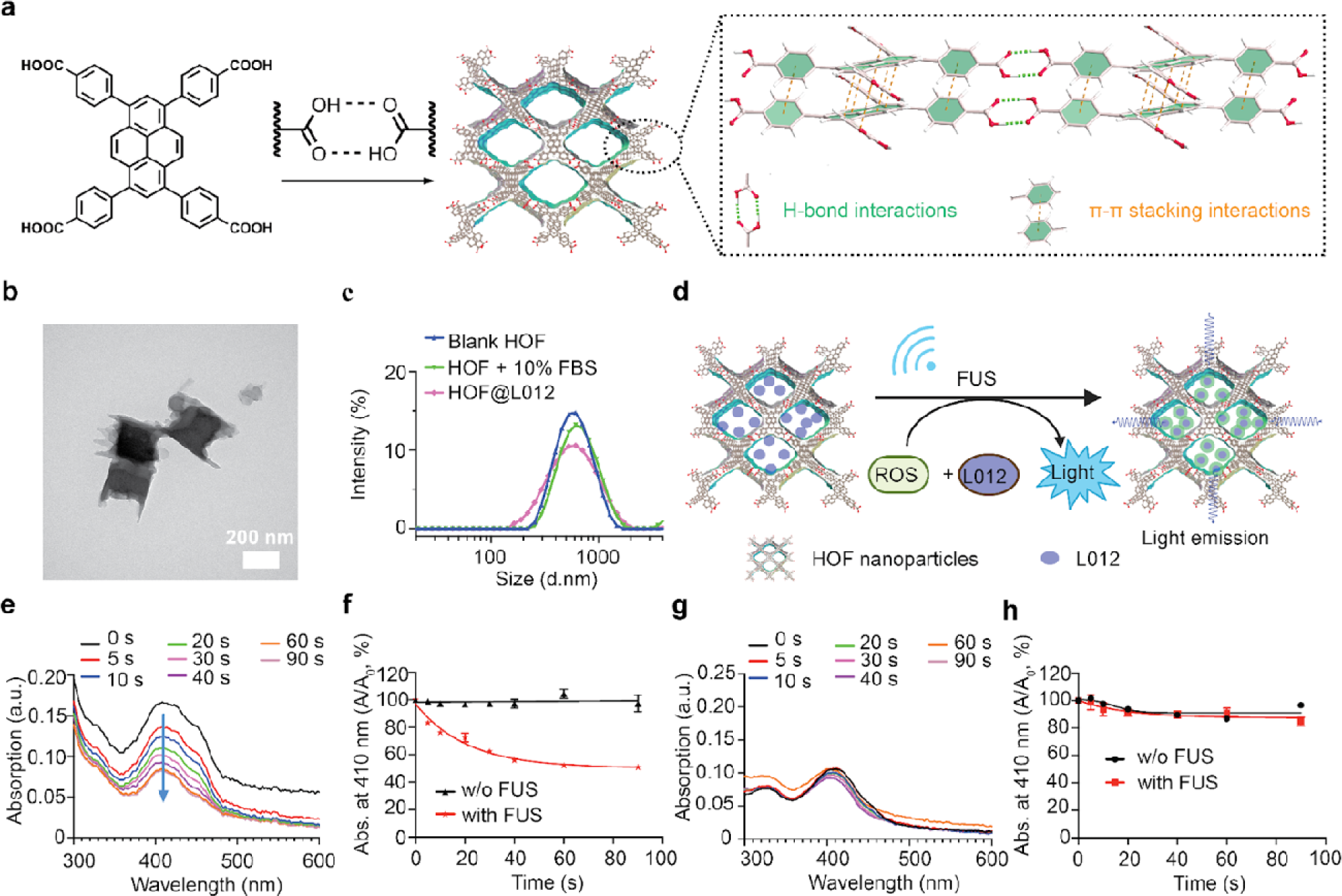
Ultrasound-triggered generation of reactive oxygen species (ROS) in HOF nanoparticles for light emission. (a) Schematic illustration of the sonosensitized HOF nanoparticles preparation. (b) The morphology of HOF nanoparticles tested via TEM, scale bar: 200 nm. (c) Hydrodynamic size distribution of HOF nanoparticles at different conditions. (d) Schematic illustration of the ultrasound-triggered cascade reaction in HOF nanoparticles. (e) The evaluation of ^1^O_2_ generation by HOF nanoparticles under ultrasound irradiation (1.5 MHz, 1.55 MPa) over time through monitoring the UV-Vis spectra of the DPBF probe. (f) The quantitative analysis of ^1^O_2_ generation through monitoring DPBF decomposition at conditions with or without ultrasound irradiation (n > 3 per group). (g) The determination of ^1^O_2_ generation in L012-loaded HOF nanoparticles and (h) the quantitative analysis of ^1^O_2_ generation.

### Sono-HOF nanoparticles act as ultrasound-triggered ROS sources for light emission

Pyrene served as an efficient photosensitizer to generate reactive oxygen species (ROS).^30^ We hypothesized that the periodic integration of H4TBAPy units into the frameworks would function as a sonosensitizer to generate ROS under ultrasound stimulation (**Fig. 1d**). We assessed and quantified ROS production using colorimetric probes, specifically 1,3-diphenylisobenzofuran (DPBF) and salicylic acid (SA), to detect singlet oxygen (^1^O_2_) and hydroxyl radical (•OH), respectively.^18,26^ During ultrasound irradiation, the characteristic UV-Vis absorption peak of DPBF at 420 nm rapidly decreased, while no changes were observed in the characteristic UV-Vis absorption peak of SA at 303 nm. These results indicated the generation of ^1^O_2_ but not •OH by the Sono-HOF nanoparticles (**Fig. 1e,f**, and **Supplementary Fig. 4**). Furthermore, the yields of ^1^O_2_ increased with higher ultrasound peak pressure and displayed ultrasound power-dependent behavior (**Supplementary Fig. 5**). We anticipated that L012 would efficiently consume the generated ^1^O_2_ to produce light, preventing any potential leakage since excessive ^1^O_2_ could damage cell membranes and lead to cell death. Consequently, we also evaluated ROS generation in L012-loaded Sono-HOF nanoparticles (HOF@L012). The results confirmed that no ^1^O_2_ leakage during ultrasound stimulation in the HOF@L012 solution (**Fig. 1g, h**). L012, a potent ROS scavenger, effectively quenched ROS for light emission, preventing normal tissue from ultrasound-induced damage.

We next evaluated the mechanoluminescent performance of HOF@L012 nanoparticles under the ultrasound stimulation. The fluorescence spectrum from HOF@L012 showed that the emission wavelength at around 470 nm, which mainly overlapped with the channelrhodopsin-2 (ChR2) for optogenetic stimulation (**Fig. 2a**).^38^ We also evaluated the real-time light emission from the HOF@L012 nanoparticles via photons recording system. Compared to luminol and dioxetane derivatives, chemiluminescent L012 demonstrated a significantly higher ROS sensitivity and reaction rate constant, which ensured the temporal light was generated once under the ultrasound stimulus.^25^ In addition, opsins activation in optogenetics usually requires temporal light control and tunable pulse frequency in neuromodulation for different neurological disease treatments, such as Alzheimer’s disease.^39^ Here, time-resolved mechanoluminescence determined that this system exhibited excellent controllability and temporal resolution, from pulse frequency 1 to 10 Hz, with less than 4 ms latency (**Fig. 2b, c** and **Supplementary Figs. 6-7**), and showed great potential in optogenetic stimulation with various frequencies. We find the light intensity is positively correlated with ultrasound peak pressure due to the energy-dependent ROS generation (**Fig. 2d** and **Supplementary Fig. 8**). In addition, HOF@L012 nanoparticles exhibit higher photon yield compared with our previously reported liposomal light source at the same ultrasound power due to the enhanced drug loading capacity of L012 (**Fig. 2d and Supplementary Table 1**). ^18^ We also assessed the long-term activation of HOF@L012 nanoparticles. As depicted in **Fig. 2e**, the decay half-time of light intensity was approximately 120 s with pulse 100 ms on and 900 ms off, demonstrating the potential for approximately 3000 repeated emissions. This is twice as high as the previous liposomal system (**Fig. 2e**). Since ultrasound energy typically travels through tissue as a propagating pressure wave, attenuating exponentially with tissue depth. As depicted in **Fig. 2f**, we could reach approximately 20 mm of tissue penetration using a 1.5 MHz ultrasound wave, with nearly 40% of the original energy effectively delivered to tissues at a depth of 10 mm in porcine skin mimic tests (**Fig. 2g** and **Supplementary Fig. 9**). This level of tissue penetration far surpasses what can be achieved with both visible and NIR light sources.^40^ Furthermore, the HOF@L012 nanoparticles demonstrate enhanced photon yields compared to the previous liposomal system at different tissue penetration depth (**Fig. 2g**), highlighting their superior potential for light delivery in deep tissue for sono-optogenetics.^18^

**Fig. 2.**
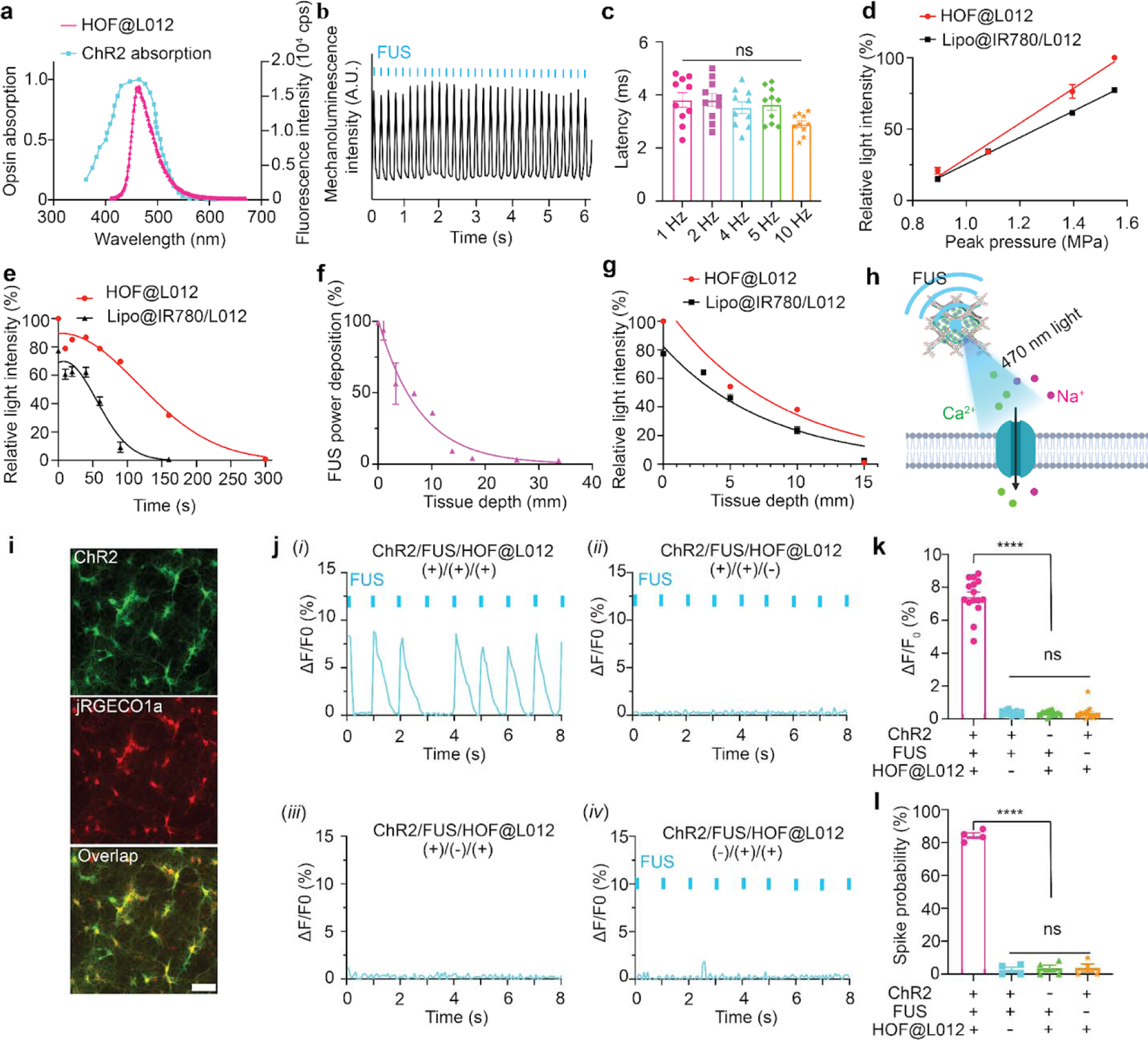
Ultrasound-triggered light emission and *in vitro* neuronal activation. (a) Mechanoluminescence spectra of HOF@L012 nanoparticles. The emission spectrum of the nanotransducers is mainly overlaid with the ChR2 opsin absorption spectrum (blue dot curve). (b) Blue light is emitted from HOF@L012 nanotransducers even at a frequency of 5 Hz under the ultrasound stimulus (1.5 MHz, 1.55 MPa, pulse 50 ms on, 150 ms off). (c) The latency between light emission and ultrasound stimulus at different frequencies. (d) Ultrasound peak pressure dependent light emission in HOF@L012 nanotransducers. (e) The light emission half-time determination of HOF@L012 and liposomal nanotransducers under the ultrasound stimulus (1.5 MHz, 1.55 MPa, pulse 1 s on 1 s off). (f) Normalized ultrasound energy deposition in the tissue and (g) normalized light emission at different tissue depths (1.5 MHz, 1.55 MPa). (h) The scheme of ChR2 opsin activation under the ultrasound activation of mechanoluminescent HOF@L012 nanoparticles. (i) Fluorescent images of mouse primary neurons expressing hSyn::ChR2-EYFP and hSyn::JRGECO1a, scale bar: 50 μm. (j) The determination of ChR2 expressing primary neurons activation through monitoring Ca^2+^ indicator (JRGECO1a) fluorescence signal changes in different experimental conditions, (*i*) FUS -, HOF@L012 -; (*ii*) FUS +, HOF@L012 -; (*iii*) FUS -, HOF@L012 +; (*iv*) FUS +, HOF@L012 +, FUS stimulation (1.5 MHz, 1.55 MPa, pulse 100 ms on 900 ms off). (k) Statistical analysis of JRGECO1a signal changes at different conditions (n=3 per group, two-way ANOVA, and multiple comparisons test). (l) Spike probability of ChR2 expressing primary neurons under the mechanoluminescene irradiation (n=3 per group, two-way ANOVA and multiple comparisons test). All plots show mean ± SEM unless otherwise mentioned. \**P* <0.05, \*\**P*<0.01, \*\*\**P*<0.001, \*\*\*\**P*<0.0001; ns, not significant.

We assessed the biosafety of HOF@L012 nanoparticles through cell viability tests and hemolysis assays. As shown in the **Supplementary Fig. 10**, the results demonstrated no significant toxicity to HEK cells, and hemolysis did not occur even at high concentrations of HOF@L012 nanoparticles, indicating the excellent biosafety. Then, we conducted *in vitro* opsin activation experiments using primary neurons under sono-optogenetics (**Fig. 2h**). Initially, neurons were transduced with AAV9-hSyn::ChR2-EYFP and AAV9-hSyn::NES-JRGECO1a to express ChR2 opsin and JRGECO1a calcium indicator (**Fig. 2i**). As shown in **Figs. 2j-l**, efficient spiking was observed in ChR2-expressing neurons, as evidenced by the tracking of calcium fluorescence signals under sono-optogenetic stimulation, resulting in an approximately 85% spike probability. In contrast, only sporadic activation was observed in calcium imaging in the absence of ultrasound stimulus or in ChR2-negative neurons.

### Sono-HOF enabled non-invasive activation of ChR2-expressing neurons *in vivo*

Next, we applied Sono-HOF enabled sono-optogenetics *in vivo* to demonstrate the intricate links between brain activity and behavior.^41^ Consequently, we examined the sono-optogenetic neuromodulation in the mouse secondary motor cortex (M2) to modulate limb motion in mice (**Fig. 3a**). Initially, we measured the light emission in the motor cortex area under ultrasound stimulation following tail vein injection. Synchronous blue light with a higher power intensity of 1.44 mW.mm^-2^ was generated in Sono-HOF system under FUS stimulation compared to the previous liposomal system (1.01 mW.mm^-2^) (**Fig. 3b**).^18^ This level of light emission was sufficient to activate ChR2 opsin for neural activation.^42^ Given that the motor cortex governs higher-order control of body movement, we monitored real-time mouse limb motions using cameras during sono-optogenetics in Thy1-ChR2-YFP transgenic mice. As depicted in **Fig. 3c-e**, DeepLabCut analysis of videos revealed that controlled limb motion was synchronized with sono-optogenetic stimulation frequency, with around 70 ms latency. Conversely, only sporadic motions were observed in the absence of FUS stimulation, Sono-HOF nanoparticle injection or in wild-type mice. In addition, we assessed neuronal activation signals in post-hoc tissue samples by examining the expression of the immediate early gene marker c-Fos. As shown in **Figs. 3f-g**, remarkable c-Fos signals were observed in the motor cortex area under sono-optogenetics experimental group, while minimum signals were found in other control groups. These results confirm the efficacy of this sono-optogenetic system in remote and spatiotemporal brain modulation in a non-invasive manner *in vivo*.

**Fig. 3.**
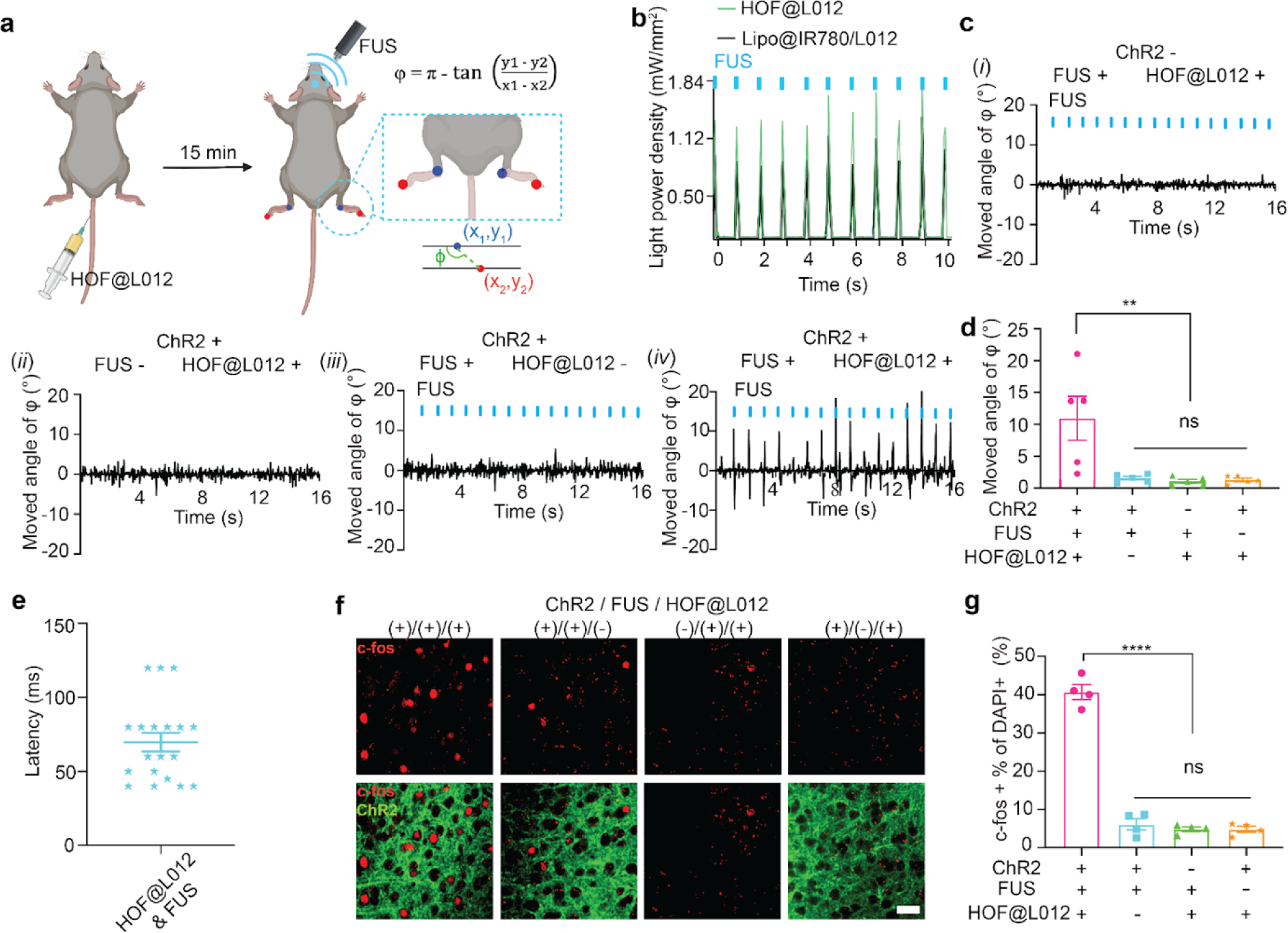
Non-invasive activation of motor cortex neurons for behavior modulation in ChR2 transgenic mice. (a) Scheme of non-invasive motor cortex stimulation via sono-optogenetics. (b) *In vivo* light generation from HOF@L012 and liposomal nanotransducers in mice motor cortex under the ultrasound stimulation (1.55 MPa, pulse 100 ms on 900 ms off). (c) Time-resolved limb’s motion tracking when the mice were treated under the different experimental conditions, (i) ChR2 -, FUS +, HOF@L012 +;(ii) ChR2 +, FUS -, HOF@L012 +; (iii) ChR2 +, FUS +, HOF@L012 - and (iv) ChR2 +, FUS +, HOF@L012 +; (d) Statistical analysis of limbs’ motions in different groups of subjects (n=5 per group, two-way ANOVA and multiple comparisons test). (e) The latency between sono-optogenetic stimulation and limb motion. (f) Confocal imaging of the expression of immediate early gene marker c-Fos in the mice motor cortex, scale bar: 30 μm. (g) Statistical analysis of c-Fos signals at different conditions at the M2 motor cortex region (n=4 per group, two-way ANOVA, and multiple comparisons test). All plots show mean ± SEM unless otherwise mentioned. \**P* <0.05, \*\**P*<0.01, \*\*\**P*<0.001, \*\*\*\**P*<0.0001; ns, not significant.

### Sono-optogenetic activation of PV-GPe neurons restores movement in hemiparkinsonian rats

A recent study has identified PV-GPe neurons as crucial components of the basal ganglia circuit, capable of inducing long-lasting attenuation of pathological activity in the SNr.^8^ Targeted interventions within the globus pallidus externa (GPe), rather than broad global interventions, have proven effective in sustaining lasting improvements in behavior, such as reducing immobility and bradykinesia, and in restoring normal physiological basal ganglia output in the 6-OHDA mouse model.^8^ Consequently, targeted activation of PV-GPe neurons through non-invasive sono-optogenetics holds great promise for ameliorating movement dysfunction in Parkinson’s disease. Here, we have selected rats due to their thicker skulls and larger brain tissue depths, making them crucial for clinical translation. We first evaluated the photon delivery efficiency of our system in the rat GPe brain area with our FUS system of focus length of 5 mm (**Supplementary Fig. 11**). As shown in **Fig. 4b**, the ultrasound triggered light intensity was around 1.22 mW.mm^-2^ in GPe after two days of intracranial injection of HOF@L012 nanoparticles. This level of light emission is expected to be sufficient to activate ChR2 opsins for neuron firing.^42^ Subsequently, we recorded neuronal excitation to sono-optogenetic stimulation within the GPe region using fiber photometry. GPe neurons were transduced with ChR2 opsins (AAV9-hSyn::ChR2-EYFP) and a red fluorescent calcium reporter (AAV9-hSyn-NES-JRGECO1a), as illustrated in **Fig. 4c**. The JRGECO1a signal exhibited a substantial increase upon ultrasound stimulation of the GPe region with ChR2 expression, while no changes were observed in the absence of ultrasound stimulation or ChR2 expression (**Fig. 4d, e**). Furthermore, we assessed neuronal activation in post-hoc tissue samples by examining the expression of the immediate early gene marker c-Fos after transducing GPe neurons with ChR2 opsins (pAAV-Ef1a-DIO hChR2(E123T/T159C)-EYFP) in PV-Cre transgenic rats. As depicted in **Fig. 4f, g**, our results revealed the specific expression of ChR2 opsins on PV-expressing GPe neurons, suggesting the feasibility for cell-type-specific modulation in deep brain stimulation. Importantly, significant c-fos signals were detected within ChR2-expressing PV-GPe neurons, whereas sporadic c-fos signals were detected in the absence of ChR2 opsins, ultrasound stimulus, or nanoparticles.

**Fig. 4.**
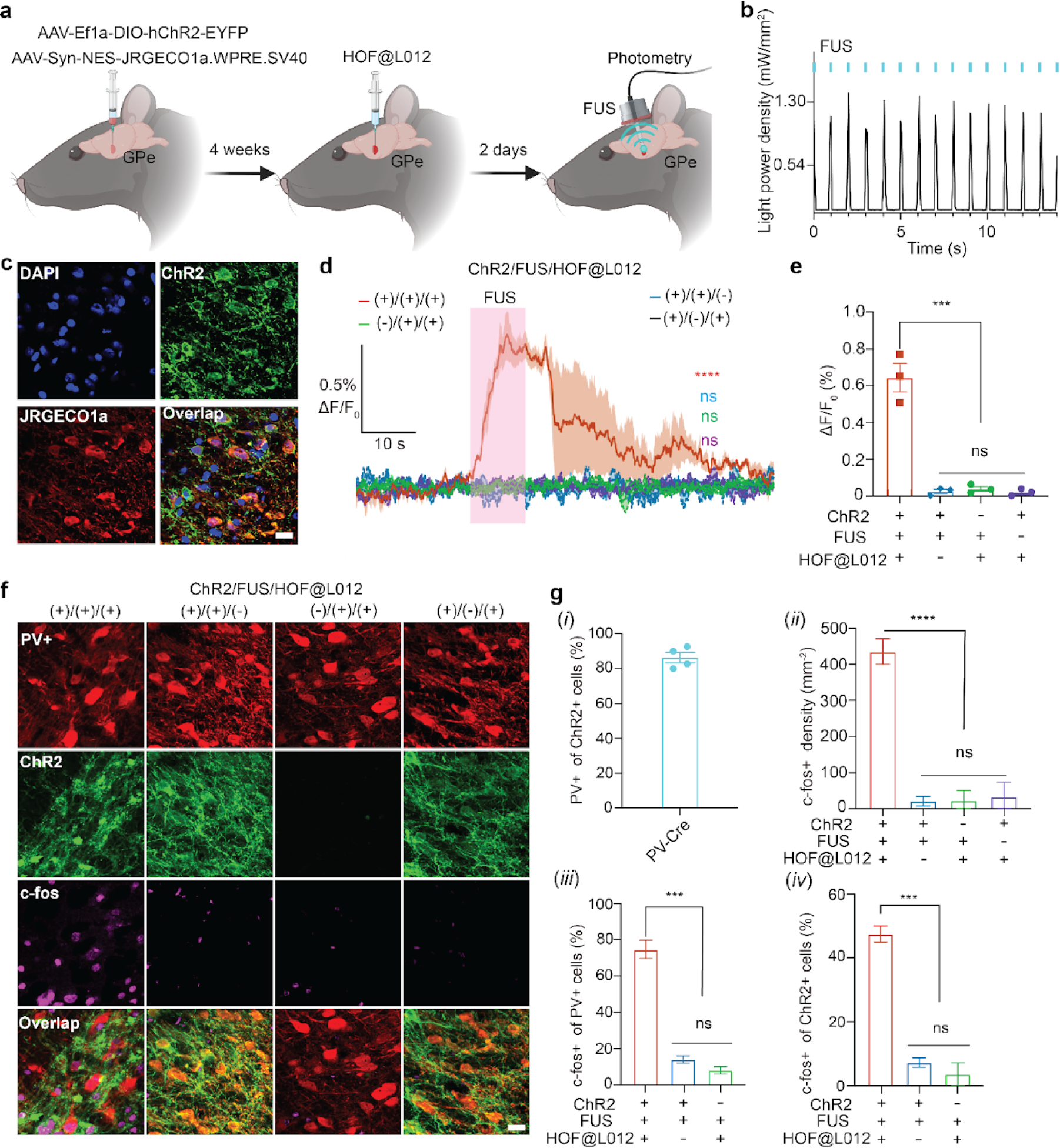
Sono-optogenetic activation of parvalbumin-expressing globus pallidus neurons in rats. (a) Experimental scheme of the *in vivo* fiber photometry of the parvalbumin-expressing globus pallidus externa (PV-GPe) neuronal activity. (b) *In vivo* light generation from HOF@L012 nanotransducers in rat GPe region under the ultrasound stimulation (2.45 MPa, pulse 100 ms on 900 ms off). (c) Confocal fluorescence images of the co-expression of ChR2 and jRGECO1a in rat GPe brain area. Scale bar: 16 μm. (d) Normalized jRGECO1a fluorescence intensity change (ΔF/F_0_) in rat GPe under the different experimental conditions. The purple patterned area represents the FUS irradiation (1.5 MHz, 1.40 MPa, pulse 10s). Solid line, mean; shade area, SEM; 3 rats in each group (n = 3). One-way ANOVA and Tukey’s multiple comparison test (P ≥ 0.05 (ns), **** P < 0.0001). (e) Statistical analysis of calcium signal changes in rat GPe region under different conditions. Mean ± SEM, n = 3 independent mouse. Two-way ANOVA and Tukey’s multiple comparison test (P ≥ 0.05 (ns), *** 0.0001≤ P < 0.001). (f) Confocal fluorescence images show the c-fos expression in the GPe after the rats are treated with different conditions. Scale bar: 20 μm. (g) (*i*) Percentage of PV-expressing neurons in ChR2-expressing GPe neurons, *(ii)* quantification of the c-fos expression cells density*, (iii)* c-fos expression percentage among PV+ neurons and *(iv)* c-fos expression percentage among ChR2+ neurons. Mean ± SEM, n = 3 independent samples. Two-way ANOVA and Tukey’s multiple comparison test (P ≥ 0.05 (ns), * 0.01≤ P < 0.05, ** 0.001≤ P < 0.01, *** 0.0001≤ P < 0.001, **** P< 0.0001).

Given the efficient activation of specific PV neurons in GPe, we next evaluated whether the sono-optogenetics could rescue the movement dysfunction in PD rats. Transgenic rats expressing Cre recombinase in PV-expressing neurons were used to build the hemiparkinsonian PD rat model. AAV9 viral vectors carrying ChR2 genes (pAAV-Ef1a-DIO hChR2(E123T/T159C)-EYFP) were intracranially injected into the GPe to target the expression of ChR2 in PV neurons (**Fig. 4f**). Subsequently, we administered the neurotoxin 6-hydroxydopamine (6-OHDA) unilaterally into the medial forebrain bundle (MFB) to induce a hemiparkinsonian model (**Fig. 5a**).^43,44^ Immunohistochemical staining confirmed a significant loss of dopamine neurons in the right substantia nigra pars compacta, validating the successful establishment of a dopamine-depleted rat model (**Supplementary Fig. 12**).^45^ After an incubation period of 4 weeks, HOF@L012 nanoparticles were injected into the right GPe region. Sono-optogenetic stimulation of PV-GPe neurons in the hemiparkinsonian rat model was conducted to validate the alleviation of movement dysfunction through the cylinder test (**Fig. 5a-c**) and the apomorphine-induced rotation test (**Fig. 5d-h**) after two days of recovery from the surgery.^45,46^ Similar to the effects of optogenetic stimulation in PV-GPe neurons, our sono-optogenetics also yielded notable improvements in movement dysfunction and increased paw usage, as evidenced by a sustained increase in contralateral touch percentages that endured for over 15 minutes following PV-GPe FUS stimulation (**Fig. 5b-c**). We also evaluated the motor recovery performance in the PD rats through apomorphine-induced rotation tests (**Fig. 5d**). As demonstrated in the **Fig. 5e-f**, PV-GPe sono-optogenetic stimulation elicited a significant reduction in the apomorphine-induced rotation rate (**Fig. 5d-f**). Motion cameras were employed to track rats’ rotation patterns during the test period, revealing decreased rotation activity in rats with sono-optogenetics (**Fig. 5e**). Additionally, dynamic motion analysis demonstrated efficient anti-rotation after sono-optogenetics, disrupting and reducing the uniform rotation speed induced by apomorphine (**Fig. 5f**). Quantification of angular speed indicated a sustained reduction in rotations over 15 minutes following PV-GPe FUS stimulation (**Fig. 5g, h**). These behavioral assessments consistently demonstrated the effectiveness of sono-optogenetics in rescuing motor dysfunction in dopamine-depleted PD rats.

**Fig. 5.**
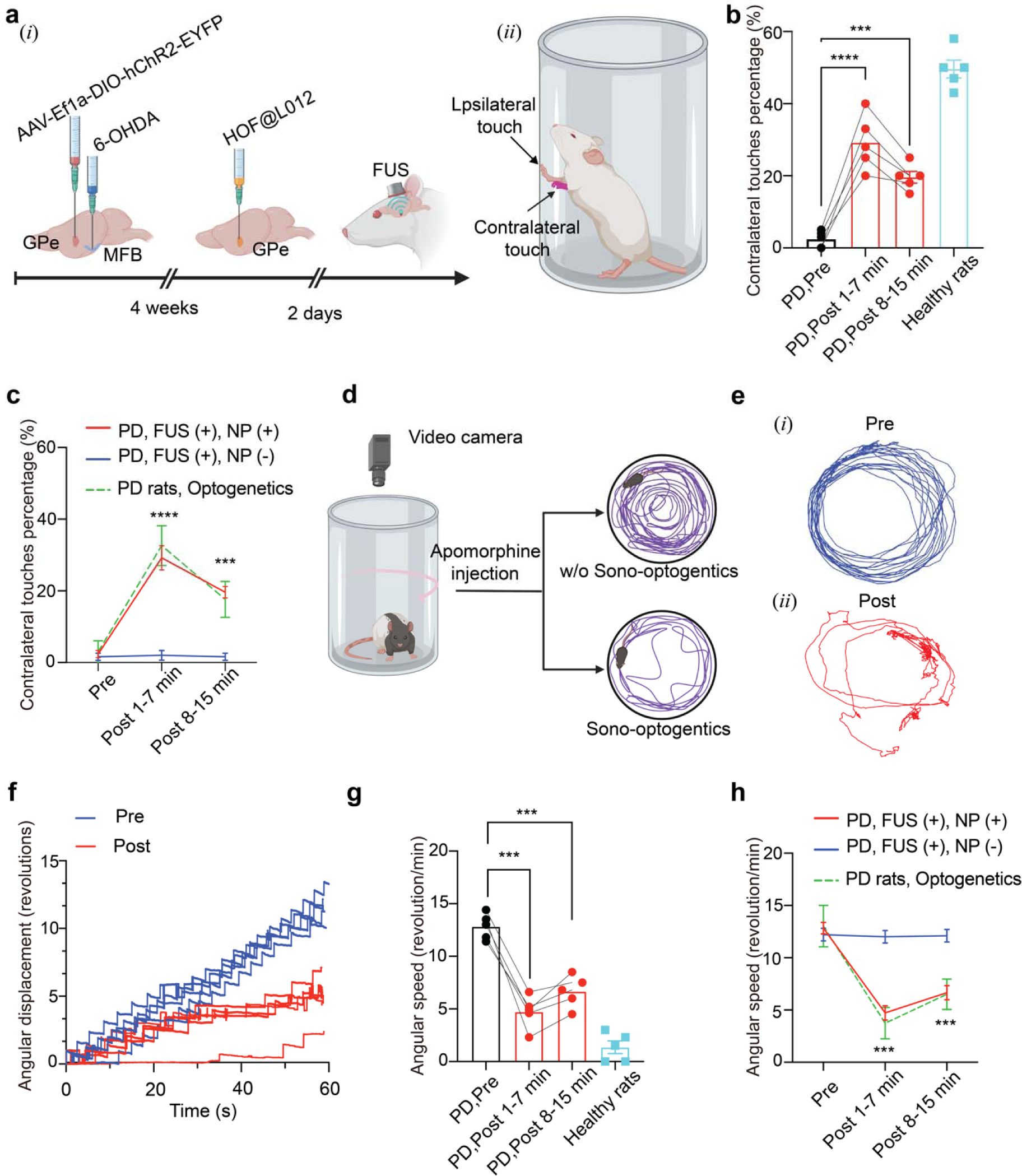
Non-invasive sono-optogenetic selective stimulation of PV-GPe neurons for long-lasting motor recovery in hemiparkinsonian rats. (a) Experimental scheme of hemiparkinsonian model creation for PV-GPe sono-optogenetics *(i)* and *(ii)* treatment efficacy assessment using cylinder test. (b) Individual contralateral touches percentage changes for each rat after FUS stimulation. Mean ± SEM, n = 5. Two-way ANOVA and Tukey’s multiple comparison test (*** 0.0001 ≤ P < 0.001, **** P < 0.0001). (c) Sono-optogenetic stimulation *(red line)* demonstrated a significant effect, comparable with optogenetics *(green dashed line)*, in terms of motor recovery of the lesioned side and an increase in contralateral touches percentage. Mean ± SEM, n = 5. Two-way ANOVA and Tukey’s multiple comparison test (*** 0.0001 ≤ P < 0.001, **** P < 0.0001). (d) Experimental scheme of PV-GPe sono-optogenetic stimulation efficacy assessment using apomorphine-induced rotation test. (e) Trace of rat during the apomorphine-induced rotation tests (i) before and (ii) after FUS stimulation. (f) Time-resolved rat rotation angular displacement during the rotation test before and after stimulation. (g) Individual angular speed changes for each rat after FUS stimulation. Mean ± SEM, n = 5. Two-way ANOVA and Tukey’s multiple comparison test (*** 0.0001 ≤ P < 0.001). (h) Sono-optogenetic stimulation *(red line)* showed a significant effect, comparable with optogenetics *(green dashed line)*, in terms of attenuation of rotation rate (angular speed). Mean ± SEM, n = 5. Two-way ANOVA and Tukey’s multiple comparison test (*** 0.0001 ≤ P < 0.001). articles and the ensuing sono-optogenetic stimulation on brain tissue integrity.

In our PV-GPe optogenetics and sono-optogenetics experiments, we achieved motor treatment effects that persisted beyond stimulation, extending up to 15 minutes in duration. However, it is worth noting that the longevity of this effect was lower compared to a previously demonstrated study conducted in a dopamine-depleted mouse model.^8^ Several factors may contribute to this discrepancy, including differences in the Parkinson’s disease models employed, the animal species used (rats vs mice), the extent of dopamine depletion, and the specific behavioral tests conducted. In the previous study, authors reported a prolonged and persistent effect during the open field test, but only in the mouse model with bilateral advanced dopamine loss.^8^ They were unable to demonstrate a persistent component of behavioral rescue in mice with partial dopamine depletion (mice with >20% striatal tyrosine hydroxylase remaining on either side). Furthermore, in unilaterally dopamine-depleted mice, PV-ChR2 stimulation did not induce enduring alterations in behavior.^8^ In our research, using a hemi-parkinsonian rat model with unilateral partial dopamine depletion, we observed a stable effect that endured beyond the cessation of PV-GPe neuronal activation. Notably, the sono-optogenetic effect closely paralleled that of optogenetics in terms of motor rescue and longevity. This suggests that sono-optogenetics could achieve a stable behavioral motor effect comparable to invasive optogenetics techniques. Finally, we also conducted a series of histological biosafety studies, encompassing Iba-1 (a marker for microglia activation), GFAP (a marker for astrocytic cells), and cleaved Caspase-3 (an apoptosis marker) staining (**Supplementary Fig. 13**) to evaluate the biosafety and immune-response of this sono-optogenetic approach, and there were no obvious activation of microglia and astrocytes and neuron apoptosis were found after two weeks sono-optogenetics. The results collectively affirmed the safety of the administered nanoparticles.

## Discussion

In this study, we have introduced the concept of sonosensitized HOF nanoparticles, loaded with chemiluminescent L012, to serve as ultrasound-triggered nano light sources. These Sono-HOF nanoparticles not only exhibited very high L012 loading capacity due to their high porosity but also acted as ROS generators, resulting in brighter and longer-lasting light emissions than previously used liposome nanoparticles. Furthermore, we extended our sono-optogenetics method for its first application in rats, specifically activating PV-GPe neurons in dopamine-depleted Parkinson’s disease rats and resulting in an alleviation of motor symptoms over time, with comparable efficacy with optogenetic stimulation in rats. The treatment longevity of our non-invasive, cell-type-targeted selective sono-optogenetic stimulation is expected to be improved further with the optimization of stimulation parameters. This approach should be further assessed in a Parkinson’s non-human primate model, with the potential to move closer to first-in-human clinical trials for Parkinson’s disease treatment. Finally, while the development of sono-mechanoluminescence materials is still in its early stages, we believe that our work opens the door for designing other non-invasive ultrasound-induced photon delivery systems for non-invasive optogenetics, gene regulation, immunotherapy, and bioimaging.^17,25,47^

## Methods

### HOF nanoparticles preparation

The HOF ligand, 1,3,6,8-Tetrakis(benzoic acid)pyrene (H4TBAPy), was generously provided by Dr. Banlin Chen’s laboratory. In a summarized procedure, 30 mg of H4TBAPy was dissolved in 3 mL of DMF at 60, followed by the gradual addition of 12 mL of distilled water under stirring (1000 rpm/min). After 15 minutes, the solution was harvested and subjected to centrifugation at 12,000 rpm (13,523 x g) for 5 minutes. The resultant yellow precipitate was collected, resuspended in acetone, and subjected to centrifugation at 12000 rpm (13523 x g) for 5 minutes. This washing step was repeated three times, followed by subsequent washes with ethanol and distilled water, each performed three times. Finally, the collected precipitate was resuspended in 3 mL of distilled water and stored at room temperature in a dark environment. This material was utilized for dynamic light scattering, UV-Vis spectrum, and TEM tests.

### Drug-loaded HOF nanoparticles preparation

L012-loaded and fluorescent dye (Rhodamine B, RB)-loaded HOF nanoparticles were synthesized. In a brief procedure, 3 mg of L012 or RB powder was introduced into 2 mL of HOF nanoparticles solution (5 mg/mL). The solution was gently agitated to facilitate the dissolution of L012 or RB. Subsequently, the solution was subjected to incubation in a water bath at 60 for 6 hours. Following this incubation, the solution was retrieved and subjected to centrifugation at 12,000 rpm (13,523 g) for 5 minutes. The resulting pellets were subjected to at least four washes with distilled water to eliminate any unloaded drugs. Subsequently, the collected pellets were resuspended in distilled water for subsequent tests.

### The generation of ^1^O_2_ under the FUS stimulation in HOF nanoparticles

1 mL of either blank HOF nanoparticles or L012-loaded HOF nanoparticles (HOF@L012) at a concentration of 0.1 mg/mL was mixed with 30 μL of a 1 mg/mL 1,3-Diphenylisobenzofuran (DPBF) methanol solution in the dark and placed in a glass vial. The mixture was then irradiated with or without focused ultrasound (FUS, 1.5 MHz, Image Guided Therapy) at 1.5 MHz. At a fixed time interval, 20 μL of the solution was extracted for UV-Vis spectrum tests. The characteristic absorption of DPBF around 420 nm was monitored and used to quantify the generation of singlet oxygen (^1^O_2_) due to the selective consumption of DPBF by ^1^O_2_. Ultrasound peak pressure was measured using a hydrophone (Onda Corporation, HGL-0200). At least three independent tests were conducted in the experiments.

### The generation of •OH under the FUS stimulation in HOF nanoparticles

Similar to the ^1^O_2_ detection, 1 mL of either blank HOF nanoparticles or HOF@L012 nanoparticle solution was mixed with 50 μL of a 1 mg/mL salicylic acid (SA) methanol solution in the dark. After being stimulated for a fixed time under ultrasound, 20 μL of the solution was extracted for UV-Vis spectrum tests. We evaluated the generation of •OH by tracking the consumption of SA using UV-Vis spectrum tests, where the characteristic absorption of SA was around 297 nm.

### The sono-mechanoluminescence spectrum evaluation of HOF@L012

2 mL HOF@L012 (2 mg/mL) were loaded into a cuvette and placed in the Fluorolog3 Fluorometer fluorescence spectrometer. The FUS transducer was positioned in contact with the cuvette. Subsequently, the solution was irradiated at a peak pressure of 1.55 MPa with a 10 s pulse. The emission was recorded using the fluorometer. The ChR2 absorption spectra were referenced from previous research and extracted using GetDataGraphDigitizer software.

### Blue light emission from the HOF@L012 nanoparticles

To assess real-time sono-mechanoluminescence, we monitored light emissions using a video camera (CS165MU1/M - Zelux® 1.6 MP Monochrome CMOS Camera, Thorlabs) in a dark environment. One milliliter of HOF@L012 (2 mg/mL) was loaded into a glass vial, which was then positioned atop an ultrasound transducer (1.5 MHz) with ultrasound gel filling. The video camera was placed in front of the glass vial to record the emissions. The solution was exposed to ultrasound stimulation with various pulse settings, including 1 Hz (50 ms pulse on, 950 ms off), 2 Hz (50 ms pulse on, 450 ms off), 4 Hz (50 ms pulse on, 200 ms off), 5 Hz (50 ms pulse on, 150 ms off), and 10 Hz (50 ms pulse on, 50 ms off), all at a peak pressure of 1.55 MPa. Alternatively, we tested different peak pressures with a 1-second pulse on and a 1-second pulse off or different pulse frequencies at a constant 1.55 MPa peak pressure. All parameters remained constant during video capture. To investigate the time delay between FUS stimulation and light emission, we simultaneously recorded the LED indicator light for the FUS pulse and the mechanoluminescence light. The time difference between these two emissions represented the latency. Data analysis was performed using ImageJ software.

### Ultrasound power deposition in the tissue

Ultrasound energy propagates through tissues in the form of waves. To assess the efficiency of ultrasound energy transmission within the tissue, pork skin was used as a surrogate for normal tissue. Pork skin samples of varying depths were positioned on the ultrasound transducer, with an ultrasound hydrophone subsequently placed behind the pork skin and filled with ultrasound gel. The primary ultrasound power was controlled by adjusting the output power, and the peak pressure behind the pork skin was detected in real-time via the hydrophone. The ultrasound transmission efficiency within the tissue was calculated by dividing the measured peak pressure by the initial peak pressure.

### In vitro neuron activation via sono-mechanoluminescence

We utilized primary cortical neurons isolated from mice in our experiments. In brief, pregnant C57BL/6 mice were euthanized when the pups reached 15.5 days of gestation. Subsequently, the pups were carefully removed from the abdominal cavity and placed in a dissection medium. After removing the meninges, the cortex was collected, subjected to a 12-minute trypsin incubation at 37°C, washed three times with the dissection medium, and homogenized by pipetting. The resulting solution was filtered using a cell strainer. The cells were then diluted in a neuron culture medium consisting of a Neurobasal medium supplemented with B-27, glutamine, and penicillin-streptomycin. Following cell counting, approximately 2 x 10^5^ cells were seeded into each well of poly-l-ornithine-coated 24-well plates (Day In Vitro 0, DIV 0). The cells were placed in an incubator with 7.5% CO2, and a glial inhibitor was added on DIV 3. On DIV 4, 0.5 μL of pAAV-hSyn-hChR2(H134R)-EYFP (Addgene viral prep # 26973-AAV9; http://n2t.net/addgene:26973; RRID:Addgene_26973) and 0.5 μL of pAAV.Syn.NES-JRGECO1a.WPRE.SV40 (Addgene viral prep # 100854-AAV9; http://n2t.net/addgene:100854; RRID:Addgene_100854) were added to each well. After an additional 5 days of incubation, ChR2 opsins (green) and the JRGECO1a calcium indicator (red) were successfully expressed in the neurons, enabling in vitro calcium imaging. To activate the neurons, glass vials filled with 2 mL of HOF@L012 nanoparticles (5 mg/mL) were positioned over the cells. Ultrasound stimulation (1.5 MHz, 1.55 MPa, pulse 100 ms on 900 ms off) was applied to generate blue light for neuron activation. The red JRGECO1a fluorescence signal was recorded using a Leica DMi8 fluorescence microscope equipped with a 20X air objective and a 30 ms exposure time in the red channel (Ex: 548-573 nm). We calculated the transient increase in red fluorescence (ΔF/F) by extracting fluorescence time-series data from neurons through manual segmentation using ImageJ software. The raw data were further processed using a custom MATLAB algorithm that detrended and normalized the fluorescent time-series data through second-order polynomial curve fits and baseline maximum fluorescent value extraction, compensating for photobleaching effects.

### In vitro biosafety evaluation, including cell viability tests and hemolysis tests. In vitro cell viability

We employed Human Embryonic Kidney 293 (HEK-293T) cells in our experiments. The 96-well plates were pre-incubated with a 10 μg/mL Poly-L-Ornithine solution before use. HEK-293T cells were seeded into the 96-well plates at a cell density of 10,000 cells per well in a complete DMEM medium. Subsequently, the cells were incubated overnight at 37°C in a 5% CO_2_ environment. After this incubation period, the culture medium was removed, and a fresh, complete medium containing HOF@L012 nanoparticles at various concentrations was added. The cells were then subjected to ultrasound treatment (1.5 MHz, 1.55 MPa, pulse 100 ms on, 900 ms off) for 20 s. Following this treatment, the cells were incubated for an additional 24 hours. After that, 10 μL of Cell-Titer Blue reagent (Promega Corporation) was added to each well, and the cells were incubated for another 4 h. Fluorescence intensity was measured using a Microplate reader (BioTek Synergy H4, 560ex/590em nm), and cell viability was calculated according to the following formula:

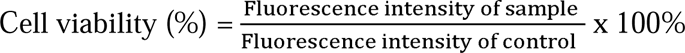

### Hemolysis tests of HOF@L012 nanoparticles

0.5 mL of fresh blood was collected from the mice’s heart. The red blood cells (RBCs) were isolated by centrifugation at 8,000 rpm for 5 minutes and subsequently washed with cold PBS at least three times. They were then suspended in 2 mL of PBS. Next, 1 mL of HOF@L012 nanoparticles at various concentrations was mixed with 20 μL of RBCs and incubated at 37°C for 2 hours. The resulting solution was then centrifuged at 12,000 rpm for 5 minutes, and the supernatant was collected for UV-Vis spectrum testing. The characteristic absorption peak at 541 nm was used to calculate the percentage of hemolysis. In addition, for control purposes, 20 μL of RBCs were added to 1 mL of distilled water to create the positive control group or 1 mL of PBS to create the negative control group. The hemolysis percentage was determined using the following formula:

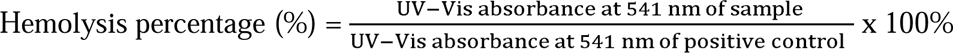

### In vivo motor cortex modulation via sono-optogenetics

We conducted our study using Thy1-ChR2-YFP transgenic mice (20-26 g; 4 weeks old; Jackson Laboratory). All procedures were designed in accordance with the National Institute of Health Guide for the Care and Use of Laboratory Animals. They were approved by the Institutional Animal Care and Use Committee at the University of Texas at Austin (AUP-2021-00086) and supported by the Animal Resources Center at the University of Texas at Austin. The mice were initially anesthetized with 2.5% isoflurane using an anesthesia machine from Vaporizer Sales & Service Inc. Their heads were securely fixed in a stereotaxic frame, and they were placed on a heating pad set to 37 to maintain body temperature. Ophthalmic ointment was applied to cover their eyes before surgery. Subsequently, their fur was shaved, and an FUS (Focused Ultrasound) transducer, filled with ultrasound gel, was positioned over the motor cortex. The coordinates of the transducer relative to bregma were as follows: anteroposterior (AP) 0.0 mm, mediolateral (ML) + 0.50 mm, and dorsoventral (DV) -0.5 mm. Next, we injected 0.2 mL of HOF@L012 nanoparticles (10 mg/mL) in saline through the tail vein. The concentration of isoflurane was then reduced to 0.5% to ensure that the mice were under light anesthesia before stimulation. To confirm the mice’s anesthesia status, we checked for any movement when gently manipulating the mice’s limbs. If there was a body movement response indicating light anesthesia, we administered ultrasound pulses (1.5 MHz, 1.55 MPa, pulse 100 ms on, 900 ms off) to activate the nanoparticles around the motor cortex, modulating limb motion. The mice’s limb movements were recorded using a video camera. Subsequently, the limb motion data were analyzed using DeepLabCut, following our previously established method.

### Stereotaxic injection of the virus into the external globus pallidus (GPe) of rats

All procedures were conducted in strict accordance with the guidelines outlined in the National Institute of Health Guide for the Care and Use of Laboratory Animals. Our research protocols received approval from the Institutional Animal Care and Use Committee at the University of Texas at Austin (Approval ID: AUP-2021-00162) and were carried out with the support of the Animal Resources Center at the University of Texas at Austin. All surgical instruments were thoroughly sterilized prior to each procedure. Before injections, the fur on the surgical area was carefully shaved, and the skin on the head underwent a meticulous sterilization process involving three rounds of cleaning with 80% ethanol and iodophor. 1,500 nL of AAV9 virus (pAAV-Ef1a-DIO hChR2(E123T/T159C)-EYFP, Addgene 35509) was injected into the GPe of Parvalbumin (PV)-Cre transgenic rats for expression of EF1a-driven, Cre-dependent, humanized channelrhodopsin E123T/T159C mutant fused to EYFP for further optogenetic activation. Virus injections were precisely administered using a micro-injection system (World Precision Instruments, UMP3 Microinjection Syringe Pump) at a rate of 300 nL/min. Following each injection, the needles were left in place inside the brain for a minimum of 5 minutes to facilitate efficient virus diffusion. They were then slowly withdrawn over a 5-minute period. Sutures were used to close the incision in the skin after the injections were completed. Following the surgical procedures, the animals were placed on a 37 heating pad and monitored in their cages until they had fully recovered.

### Transgenic Rat Breeding and Genotyping

At first, male Parvalbumin (PV)-Cre transgenic rats (the gift from the Loren Frank lab, UCSF) were bred with wild-type Long Evans female rats (Charles River Laboratories). Tissue samples from rat pups were collected using an ear-punching device and genotyped to identify PV-Cre transgenic animals (Transnetyx genotyping service), which were further used in experiments.

### Rat model for photometry tests

Wild-type Long Evans rats (3-4 months old, Charles River) were used in our experiments. Rats were anesthetized with 5% isoflurane and received a subcutaneous injection of meloxicam (2 mg/kg) and Ethiqa (0.65 mg/kg) before surgery, respectively. 1,500 nL AAV9-hSyn::ChR2-EYFP and pAAV.Syn.NES-JRGECO1a.WPRE.SV40 mixture (1:1, v/v) was unilaterally injected into the external globus pallidus (GPe), with the coordinates relative to bregma: anteroposterior (AP) -0.9 mm, mediolateral (ML) +3.00 mm, and dorsoventral (DV) -5.30 mm.

### Parkinson’s Disease Rat Model Creation

To induce Parkinson’s disease in PV-Cre rats, a unilateral hemiparkinsonian rat model was created through stereotaxic injection of 6-hydroxydopamine (6-OHDA, Hello Bio, HB1889) into the medial forebrain bundle (MFB) region.^43,44^ Under isoflurane anesthesia, a unilateral incision was made, exposing the skull. A hole was drilled unilaterally, and 2 μl of 4 μg/μl 6-OHDA in 0.9% saline was injected into the MFB (-4.0 mm AP, +1.2 mm ML, -8.1 mm DV) using a microliter syringe with a 33G needle. The infusion rate was controlled by a digital micro-syringe pump at 0.25 μl/min. The micro-syringe was left in place for an additional 5-7 minutes to ensure proper diffusion of the solution. The hemiparkinsonian symptoms were observed in 2-3 weeks after the 6-OHDA injection, and the hemiparkinsonian model was confirmed through behavioral tests (cylinder test and apomorphine-induced rotation test) and tyrosine hydroxylase (TH) immunofluorescence staining in the rat’s basal ganglia. Detailed information on the antibodies used in this work can be found in **Supplementary Table 2**.

### Photometry tests to record the neuron activation under sono-optogenetics

After a 4-week period following virus injection, the rats were utilized for photometry tests. Similar to the virus injection procedures, 2 μL of HOF@L012 nanoparticles at a concentration of 100 mg/mL were unilaterally injected into the GPe using coordinates of (-0.9 mm AP, +3.0 mm ML, -5.3 mm DV). Subsequently, optical fibers (200 μm core, R-FOC-BF200c-39NA, sourced from RWD Life Science) were implanted with corresponding coordinates in the GPe region. Following a recovery period of 2 days, the rat was restrained, and its head was immobilized. A FUS transducer (1.5 MHz, 2.45 MPa) was positioned above the head and filled with ultrasound gel. A pulse sequence (100 ms on, 100 ms off, with a duration of 10 seconds) was administered to irradiate the GPe area according to predefined FUS parameters. The resulting signal was recorded and analyzed using the R810 Dual Color Multichannel Fiber Photometry System from RWD Life Science.

### Behavior tests in PD rats under Sono-optogenetics

Cylinder and apomorphine-induced rotation tests were performed in four rat groups: naive *(n=5)*, hemiparkinsonian with PV-GP optogenetic stimulation *(n=5)*, hemiparkinsonian with PV-GP sono-optogenetic stimulation *(n=5),* hemiparkinsonian without stimulation.

### Cylinder test during optogenetic and sono-optogenetic stimulations

A cylinder test is traditionally used to measure asymmetric forelimb use in hemi-parkinsonian rats, with the extent of asymmetry indicating the severity of the unilateral lesion induced by the 6-OHDA injection.^45^ Each rat was placed in a cylindrical environment (inner diameter: 20 cm, height: 30 cm) and allowed to behave spontaneously while a video camera was positioned directly above the cylinder. Rats were recorded for five minutes with no prior habituation to the cylinder, and the number of wall touches with each paw was analyzed from the resulting videos. The data were then analyzed to calculate contralateral touch percentages for each rat, represented as the number of contralateral touches over the sum of contralateral and ipsilateral touches, multiplied by 100.

Next, we focused on PV-GPe optogenetic stimulation during the cylinder test. We connected the LED Driver (Thorlabs, LEDD1B, and M405F1) to the implanted fiber optic cannula using a patch cable. For optogenetic stimulation, various parameters were tested for motor recovery efficacy: frequencies of 5-20 Hz, pulse widths of 20-40 ms, power levels of 5-7 mW, and stimulation periods of 30-60 seconds with a 30-180 second pause. The most effective parameters for the cylinder test were found to be 7 mW, 20 Hz, with either a 15 ms on / 35 ms off pattern or a 20 ms on / 30 ms off pattern, with a 30-second stimulation period and a 30-60 second interpulse interval, resulting in a total stimulation period of 7 minutes.

Moving on, we explored PV-GPe sono-optogenetic stimulation. Similar to the virus injection procedures, we unilaterally injected 2 μL of HOF@L012 nanoparticles at a concentration of 100 mg/mL into the GPe using coordinates of (-0.9 mm AP, +3.0 mm ML, -5.3 mm DV). Then, on the 2nd and 3rd days after nanoparticle injection, we performed FUS stimulation (sono-optogenetics) of PV-GPe neurons (5-10 Hz frequency, 60-100 ms pulse width, 30-90 sec stimulation period). To achieve effective FUS stimulation while the rat was awake, we securely placed the rat in a medical-grade plastic bag with holes for the nose and the top of the head. This approach minimized stress for the rat and ensured proper positioning of the FUS transducer at the target stimulation site. After a 30-90 second FUS stimulation period, we placed the rat back into the cylinder to record wall touches. The contralateral touches percentages were calculated for each group as described above.

### Apomorphine-induced rotation test during optogenetic and sono-optogenetic stimulations

The apomorphine rotation test is commonly used to assess the extent of motor impairment induced by the lesion.^45,46^ Apomorphine, as a dopamine receptor agonist, acts post-synaptically and, due to the hyperstimulation of supersensitive dopamine receptors in the denervated striatum, induces rotation in the opposite contralateral direction. To induce rotation, we injected subcutaneously 0.1 mg/kg of apomorphine dissolved in a sodium chloride solution. After the injection, the rat was placed inside a glass cylinder (inner diameter: 25 cm, height: 50 cm). The rats typically began to rotate within about 5-10 minutes, reaching their maximum rotation rate in approximately 15 minutes. The effects of apomorphine lasted for 60-65 minutes (confirmed in the control hemiparkinsonian group). Once the maximum rotation rate was achieved (maintained for a constant 5-minute period), we recorded the baseline rate for 5 minutes. Afterward, we performed optogenetic or sono-optogenetic stimulation as described above. Optogenetic stimulation was conducted for 7 minutes, and the post-stimulation period was recorded for 15 minutes. For sono-optogenetics, video recordings were made for 20 minutes after stimulation.

Angular speed (revolutions per minute) was calculated for each group. Additionally, video recordings were processed using DeepLabCut software to track rotations within different groups.

### C-fos staining in the mice/rat brain section after Sono-optogenetics

The animals initially underwent sono-optogenetic treatment. After 60 minutes, the animals were anesthetized intraperitoneally using ketamine (16 mg/kg) and subsequently perfused with cold PBS, followed by 4% paraformaldehyde. The brains were carefully extracted and immersed in 4% paraformaldehyde at 4, where they were left to soak overnight. The brains were then sectioned into 60 μm thick slices using a vibrating blade microtome (Leica VT1200). These brain slices were rinsed with a 0.3% Triton-X PBS (TBS) solution and subsequently subjected to a 30-minute blocking step with a 5% bovine serum albumin TBS solution at room temperature. In the case of mouse brain sections, after the 30-minute blocking, the slices were incubated with a rabbit anti-c-Fos antibody (ab222699, Abcam, 1:500) in TBS. The samples were left to incubate at 4°C overnight and underwent three subsequent washes with TBS solution. Following this, a mixture of TBS and secondary antibodies (goat anti-rabbit Alexa Fluor 594, ab175652, Abcam, 1:500) was added, and the slices were incubated for 2 hours at room temperature in a dark environment. The slices were then washed three times with TBS, mounted on slides using mounting media (9990402, Fisher Scientific), and covered with coverslips. Confocal images were acquired using a Zeiss 710 laser scanning microscope.

For rat brain sections, the procedure was essentially the same, except for the choice of antibodies. After the initial 30-minute blocking step at room temperature using a 5% bovine serum albumin TBS solution, the samples were incubated with a rabbit anti-c-Fos antibody (ab289723, Abcam, 1:500) or a mouse anti-Parvalbumin antibody (P3088-100UL, Sigma-Aldrich, 1:1,000) in TBS. Subsequently, a mixture of TBS and secondary antibodies (goat anti-rabbit Alexa Fluor 647, A32733, Fisher Scientific, 1:500) or (goat anti-mouse Alexa Fluor 594, A21125, Invitrogen, 1:1,000) was applied, and the slices were again incubated for 2 hours at room temperature in a dark environment. After three washes with TBS, the slices were mounted on slides using mounting media (9990402, Fisher Scientific) and covered with coverslips for confocal imaging. Detailed information on the antibodies used in this work can be found in **Supplementary Table 2**.

### Biosafety evaluation via immunostaining

The rats initially underwent sono-optogenetic treatment. After 14 days, the rats were anesthetized intraperitoneally using ketamine (16 mg/kg) and subsequently perfused with cold PBS, followed by 4% paraformaldehyde. The brains were carefully extracted and immersed in 4% paraformaldehyde at 4, where they were left to soak overnight. The brains were then sectioned into 60 μm thick slices using a vibrating blade microtome (Leica VT1200). These brain slices were rinsed with a 0.3% Triton-X PBS (TBS) solution and subsequently subjected to a 30-minute blocking step with a 5% bovine serum albumin TBS solution at room temperature. After that, the samples were incubated with a rabbit anti-Iba1 antibody (013-27691, FUJIFILM Wako Chemical, 1:1,000) or rabbit anti-GFAP antibody (13-0300, Invitrogen, 1:1,000) or rabbit anti-Cleaved Caspase-3 (9661, Cell signaling Technology, 1:1,000) in TBS. The samples were left to incubate at 4°C overnight and underwent three subsequent washes with TBS solution. Following this, a mixture of TBS and secondary antibodies (goat anti-rabbit Alexa Fluor 594, ab175652, Abcam, 1:500) was added, and the slices were incubated for 2 hours at room temperature in a dark environment. The slices were then washed three times with TBS, mounted on slides using mounting media (9990402, Fisher Scientific), and covered with coverslips. Confocal images were acquired using a Zeiss 710 laser scanning microscope. Detailed information on the antibodies used in this work can be found in **Supplementary Table 2**.

## Acknowledgment

TEM image acquisition was performed with the help of Michelle Mikesh at the Center for Biomedical Research Support Microscopy and Imaging Facility at UT Austin (RRID# SCR_021756). Prof. Huiliang Wang acknowledges funding support from the American Parkinson Disease Association (APDA) grant, NIH Maximizing Investigators’ Research Award (National Institute of General Medical Sciences 1R35GM147408), the University of Texas at Austin Startup Fund, Robert A. Welsh Foundation Grant (No. F-2084-20210327) and Craig H. Neilsen Foundation Pilot Research Grant. We acknowledge BioRender.com for the figures drawing.

## Author Contributions

W.W., I.P., and Prof. H.W. designed the project. W.W. led this project and performed all the characterization, including materials characterization, cell tests, mice tests, data analysis and also rat tests with I.P.. Then, I.P. built the rat model, and conducted the rat behavior tests and relative immunohistology tests. Y.S. Y. X. and Prof. B.C. provided the HOFs building units and conducted NMR, XRD and gas absorption tests of HOFs. K.T. T.W., and A.S. helped with animal behavior data analysis. X.L., W.H., J.J., J.H., A.R.L., and B. A. helped in immunohistology tests, cell culture and analysis. Prof. L. F., and Prof. S. S helped with data analysis and manuscript writing. All the authors contributed to the writing of the manuscript.

## Additional Information

Supplementary Information is available for this paper.

