## Supplementary Information for "Hydrogen-bonded organic framework nanotransducers enabled sono-optogenetics for Parkinsonian rats"

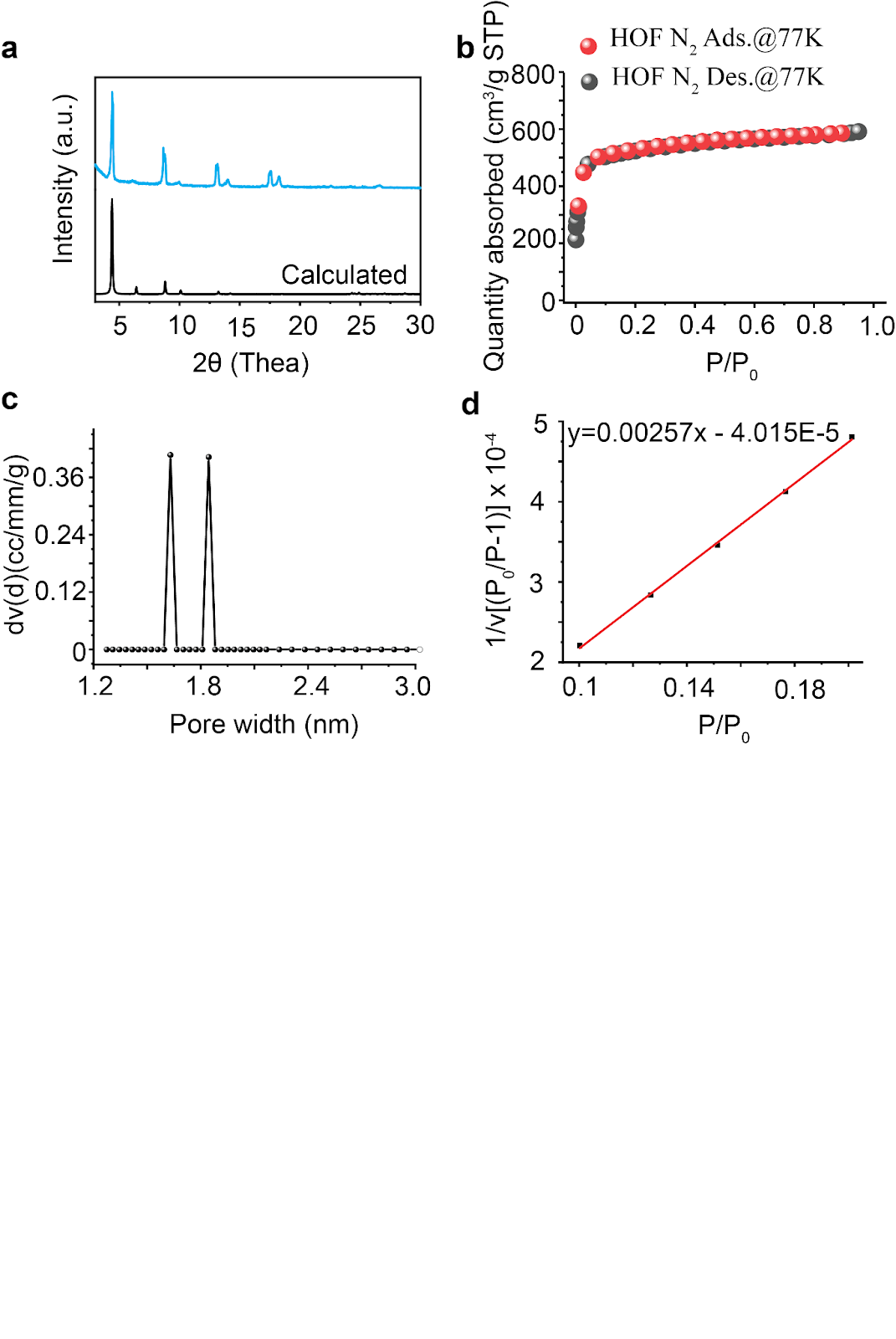

**Supplementary Fig. 1. The crystal structure determinations of HOF nanoparticles.** (a) Powder X-ray diffraction (PXRD) tests of HOF nanoparticles, where the as-synthesized HOF nanoparticles were phase-pure and highly crystalline, which fitted very well with the calculated one. (b) N_2_ sorption isotherm (adsorption and desorption) at 77 K for HOF nanoparticles, where the pore volume of HOF nanoparticles is 34.7 cm^3^mol^-1^. (c) Pore size distribution of HOF nanoparticles, and the dominant pore size is around 1.63 nm and 1.84 nm. (d) The BET surface areas for as-synthesized HOF nanoparticles (497.5 m^2^g^-1^) were obtained from the N_2_ adsorption isotherm at 77 K.

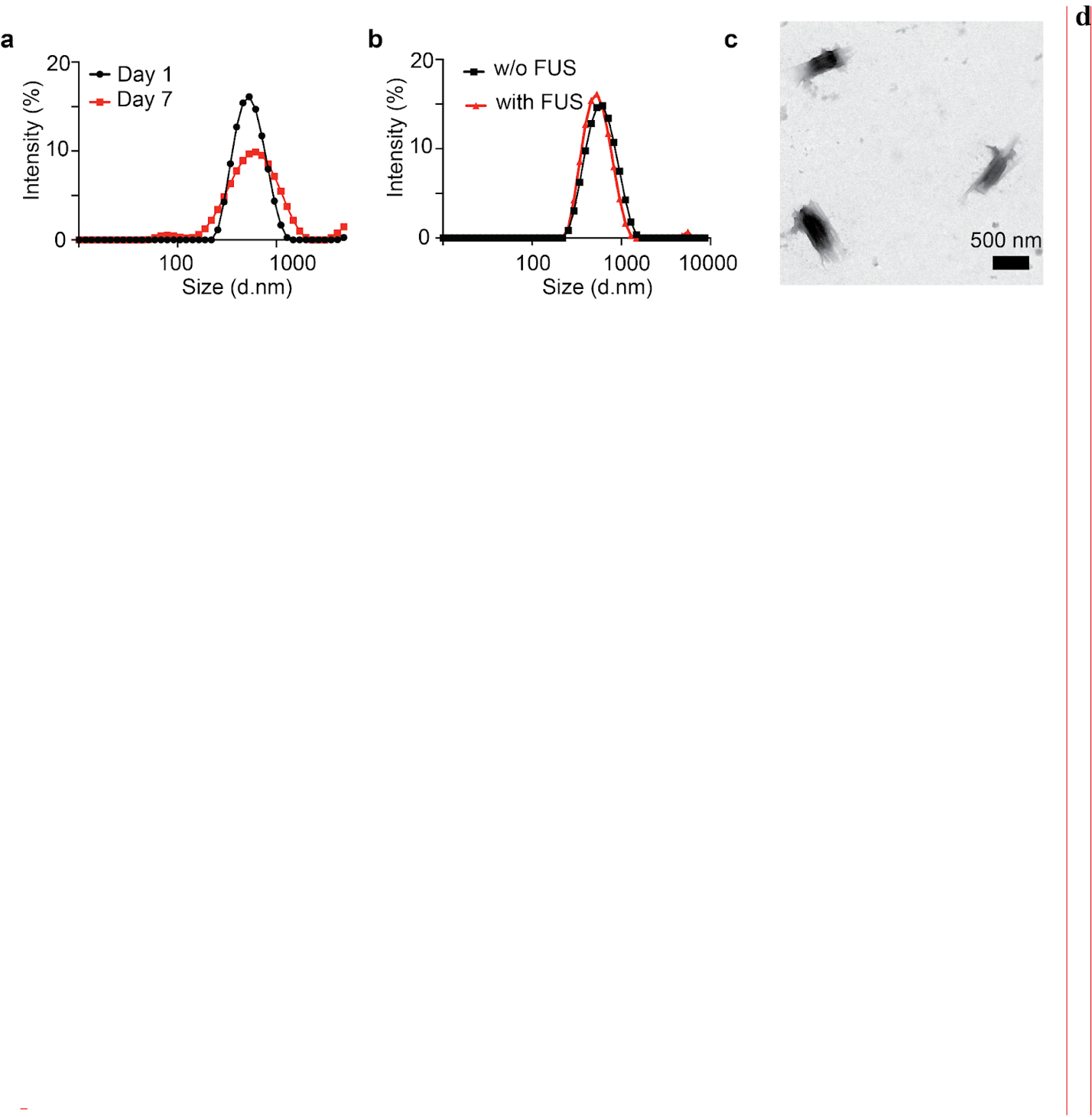

**Supplementary Fig. 2. The stability evaluation of HOF nanoparticles.** (a) The dynamic light scattering tests of HOF nanoparticles after 1 day or 7 days storage, (b) The dynamic light scattering tests of HOF nanoparticles stimulated with and without ultrasound for 60 s (1.55 MPa, 1.5 MHz). (c) The TEM images of HOF nanoparticles after 60 s ultrasound stimulation (1.55 MPa, 1.5 MHz), scale bar: 500 nm.

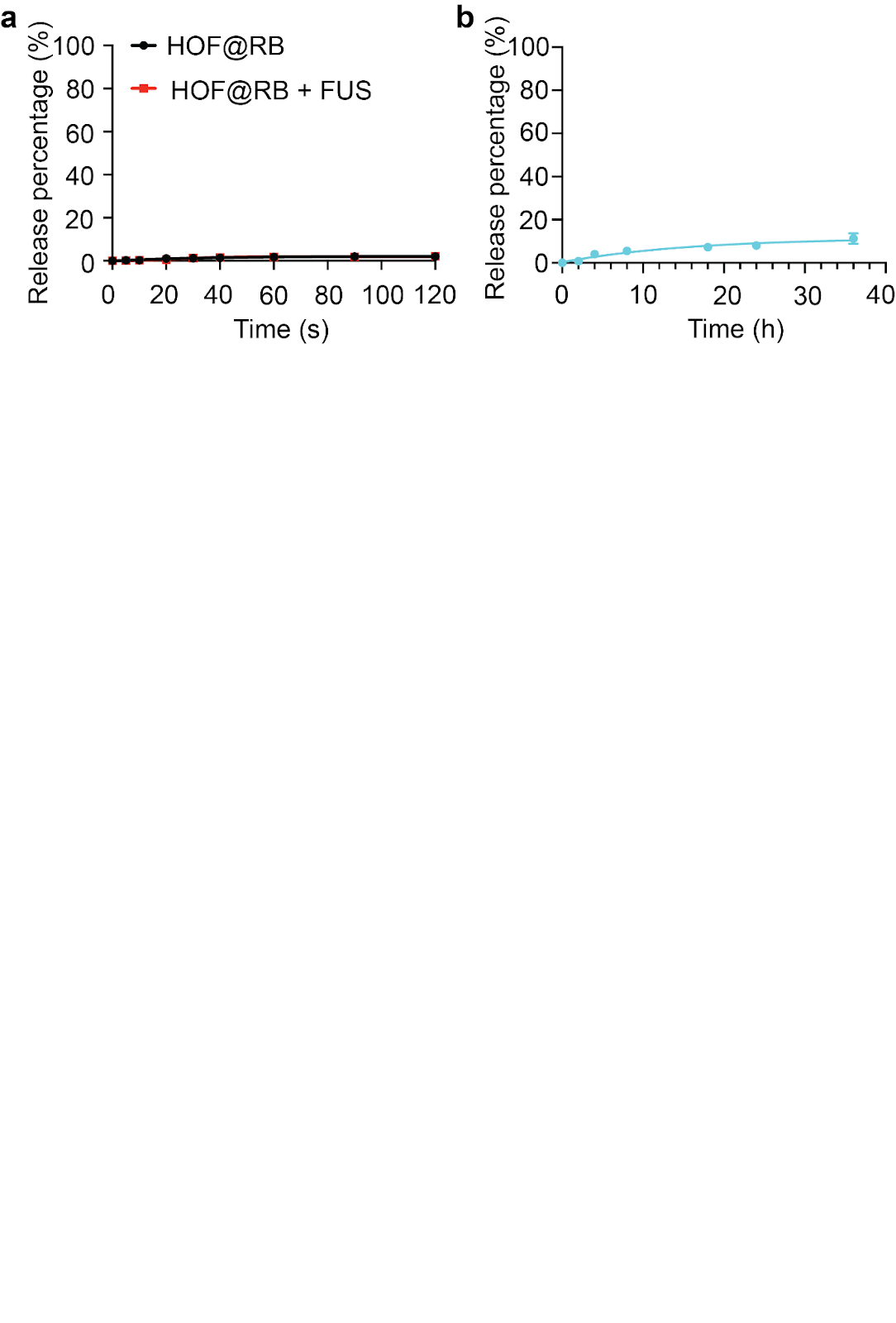

**Supplementary Fig. 3. The stability evaluation of dye-loaded HOF nanoparticles.** (a) The dye release from HOF nanoparticles under ultrasound stimulation. (b) The long-term stability evaluation of dye-loaded HOF nanoparticles, where only around 8% free due was released after incubating 36 h at 37 ℃.

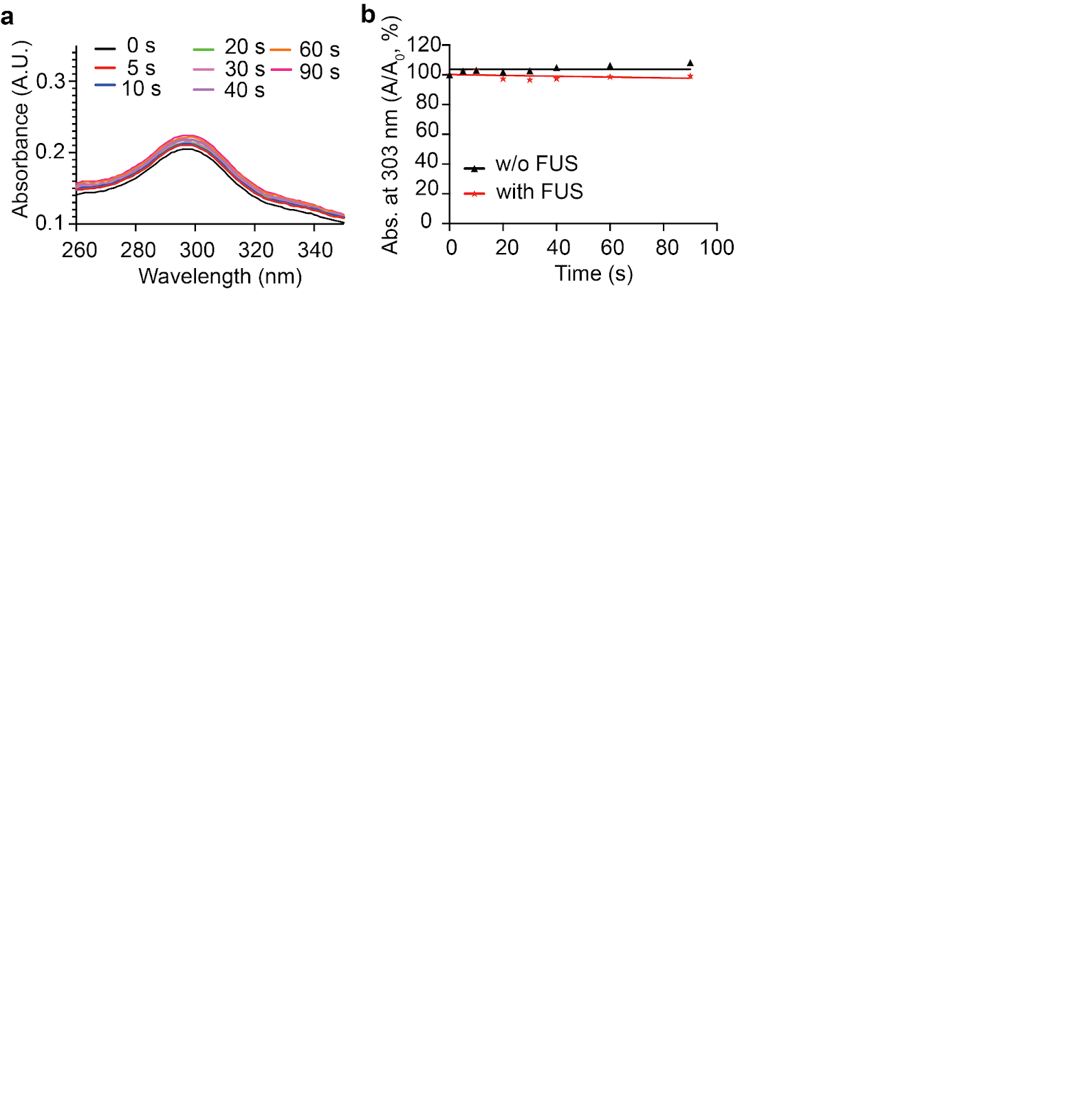

**Supplementary Fig. 4. The evaluation of the hydroxyl radicals (∙OH) generation through HOF nanoparticles under ultrasound irradiation.** (a) The evaluation of hydroxyl radicals (∙OH) generation by HOF nanoparticles under ultrasound irradiation (1.5 MHz, 1.55 MPa) over time through monitoring the UV-Vis spectra of the SA probe. (e) The quantitative analysis of ∙OH generation through monitoring SA decomposition at conditions with and without ultrasound irradiation (n > 3 per group).

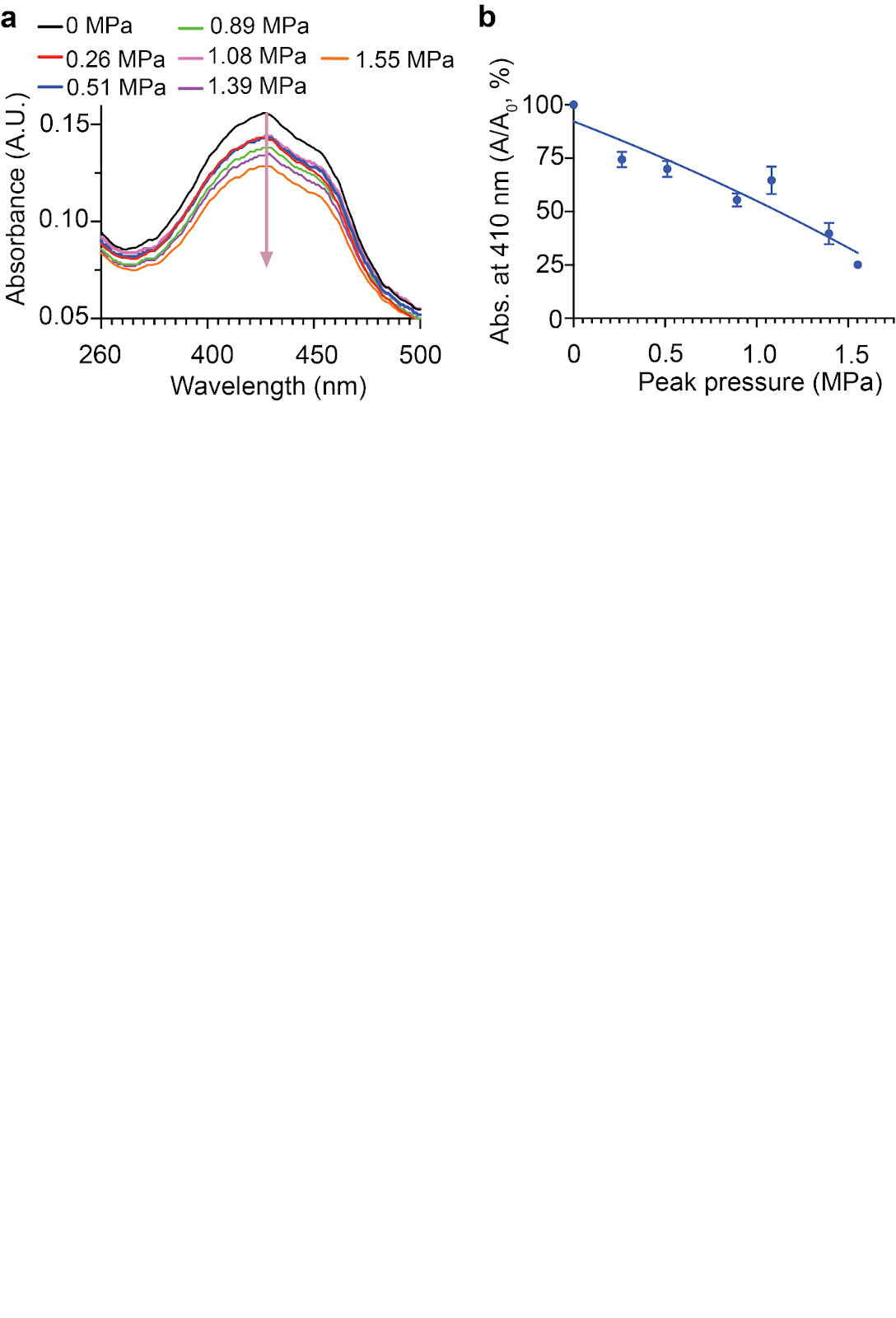

**Supplementary Fig. 5. Ultrasound peak pressure dependent ^1^O_2_ generation by HOF nanoparticles under ultrasound irradiation.** (a) The UV-Vis spectra of the DPBF probe after the HOF nanoparticles were irradiated by ultrasound for 60 s at different peak pressures. (b) The quantitative analysis of ^1^O_2_ generation through monitoring DPBF decomposition at conditions with or without ultrasound irradiation (n > 3 per group).

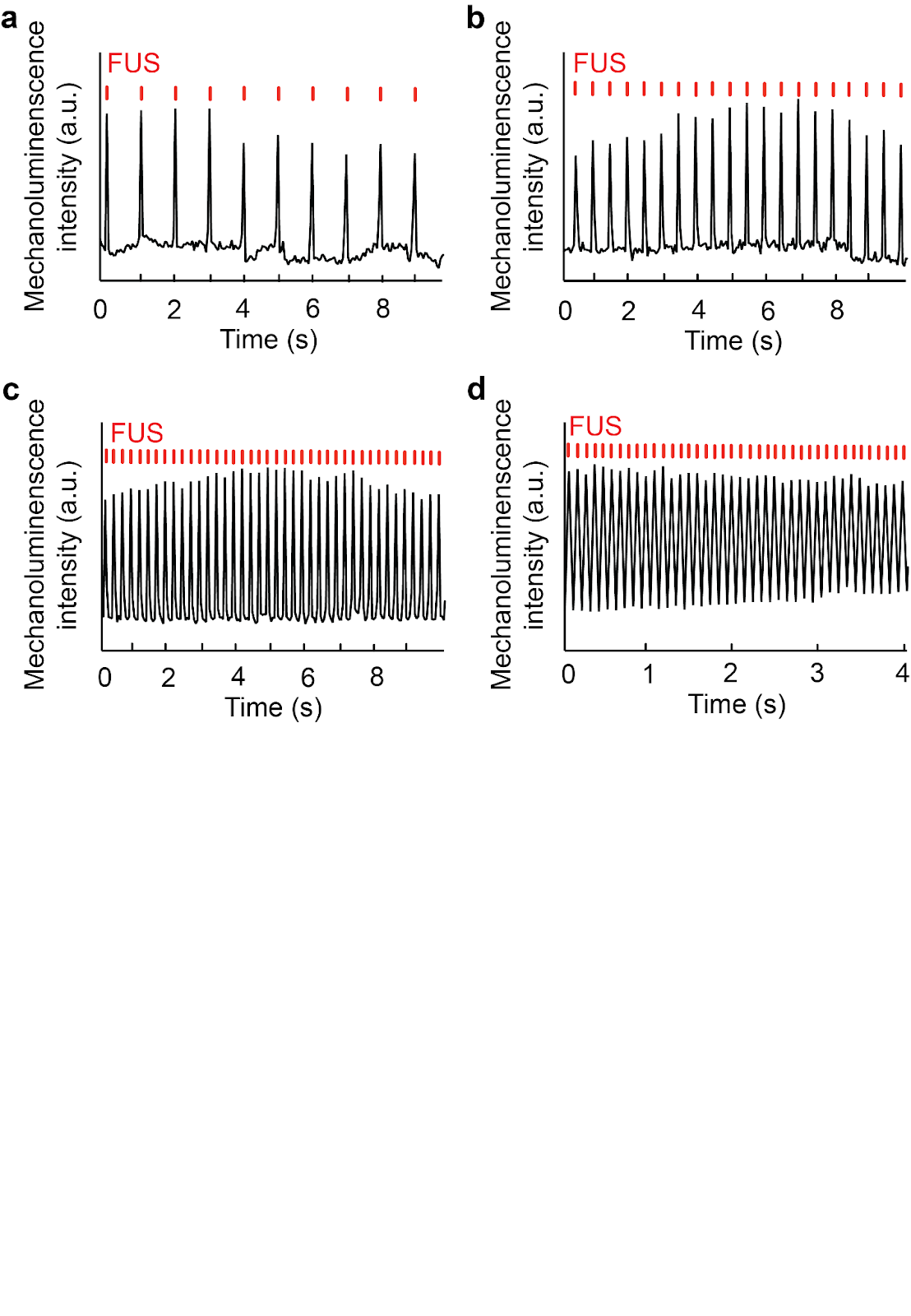

**Supplementary Fig. 6. Ultrasound triggered light emission from HOF@L012 nanoparticles at different pulse frequencies (1.55 MPa,1.5 MHz, pulse 50 ms on).** (a) 1 Hz, (b) 2 Hz, (c) 4 Hz and (d) 10 Hz.

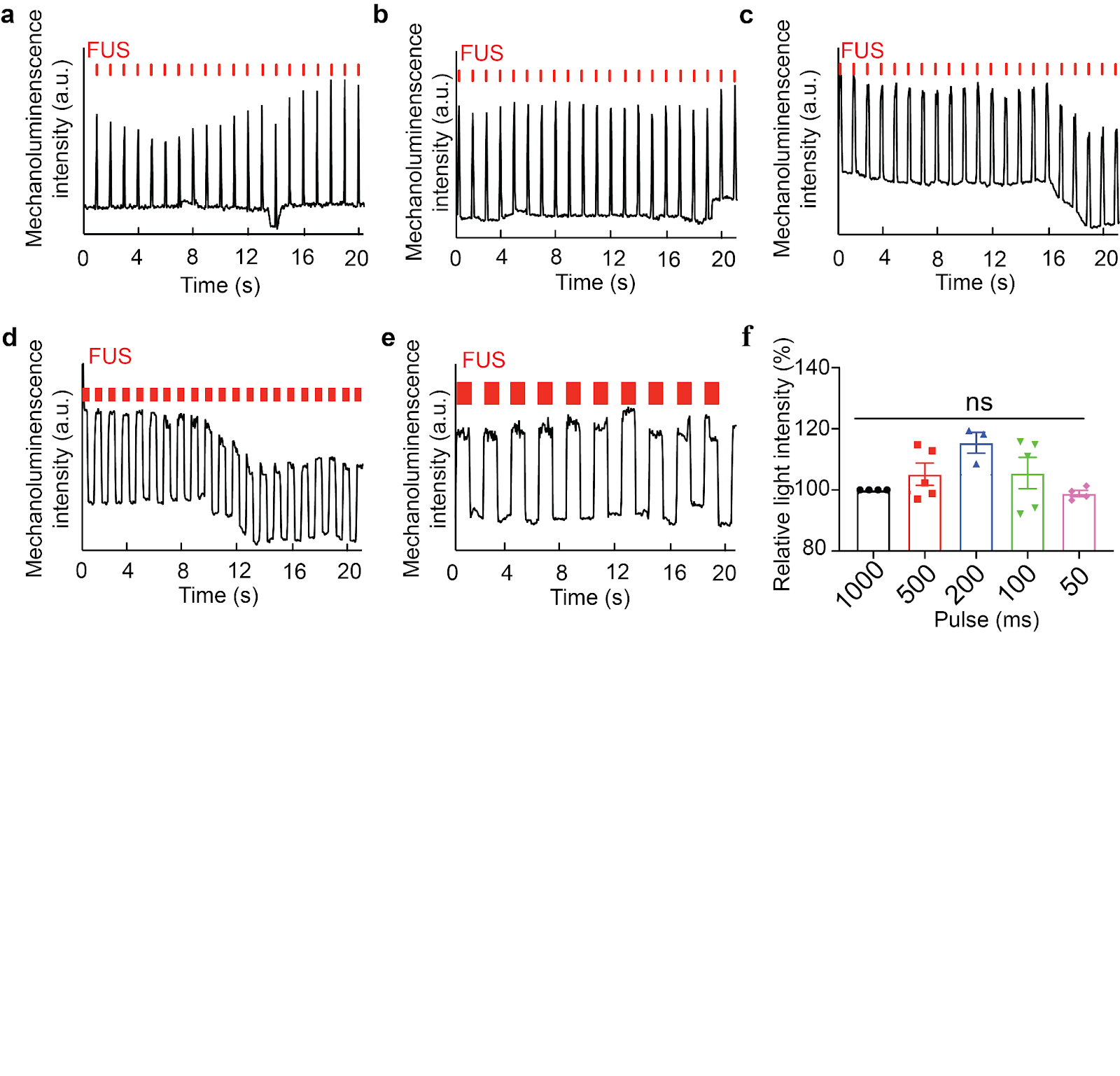

**Supplementary Fig. 7. Ultrasound triggered light emission from HOF@L012 nanoparticles at different pulse parameters (1.55 MPa, 1.5 MHz).** (a) 50 ms on 950 ms off, (b) 100 ms on 900 ms off, (c) 200 ms on 800 ms off, (d) 500 ms on 500 ms off, (e) 1000 ms on 1000 ms off, (f) statistical analysis of light intensity at different pulse parameters.

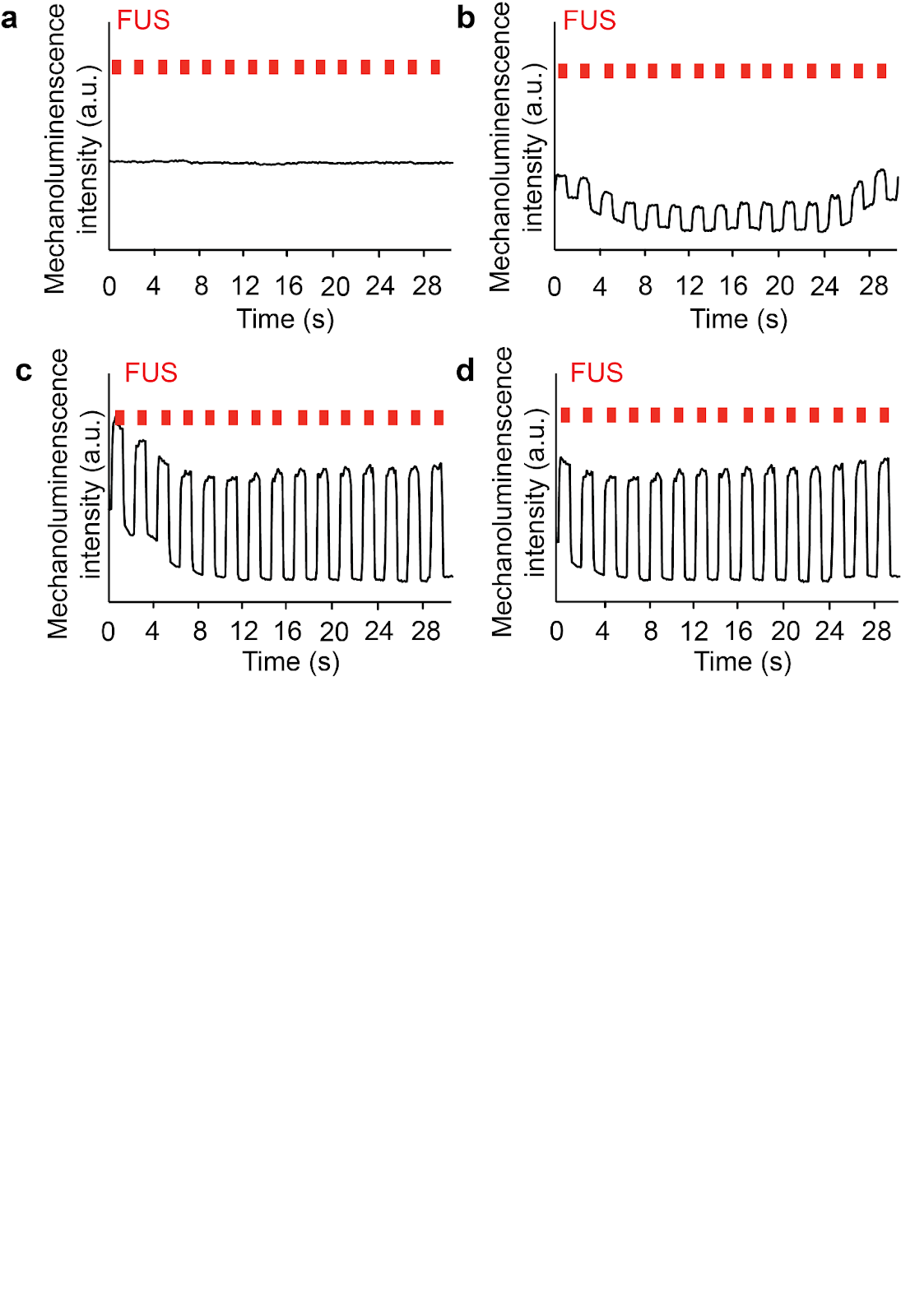

**Supplementary Fig. 8. Ultrasound triggered light emission from HOF@L012 nanoparticles at different peak pressures.** (a) 0.89 MPa. (b) 1.08 MPa. (c) 1.40 MPa. (d) 1.55 MPa.

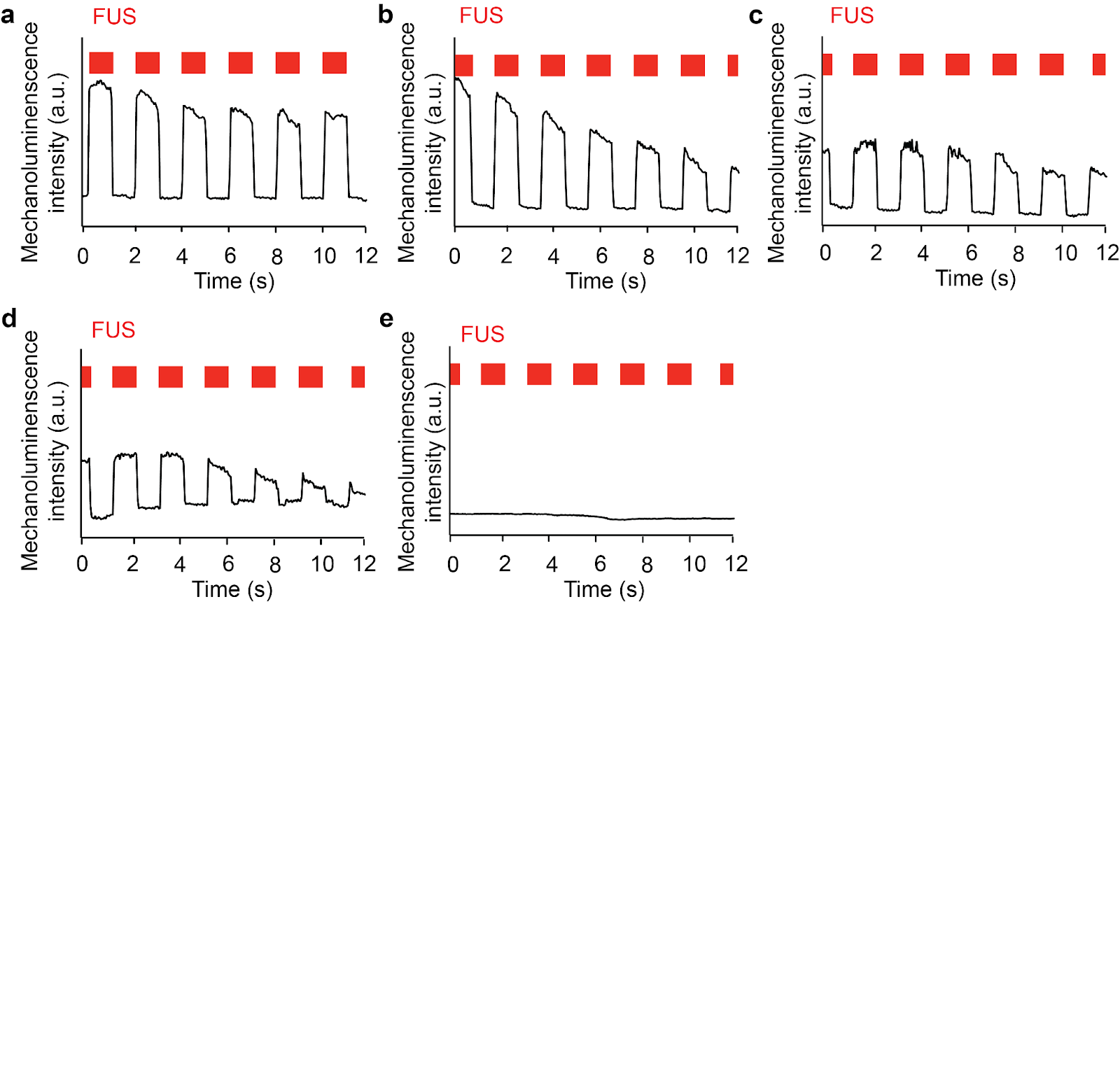

**Supplementary Fig. 9. Ultrasound triggered light emission from HOF@L012 nanoparticles at different pork skin depths**. (a) 0 mm, (b) 3mm, (c) 5 mm, (d) 10 mm, and (e) 15 mm.

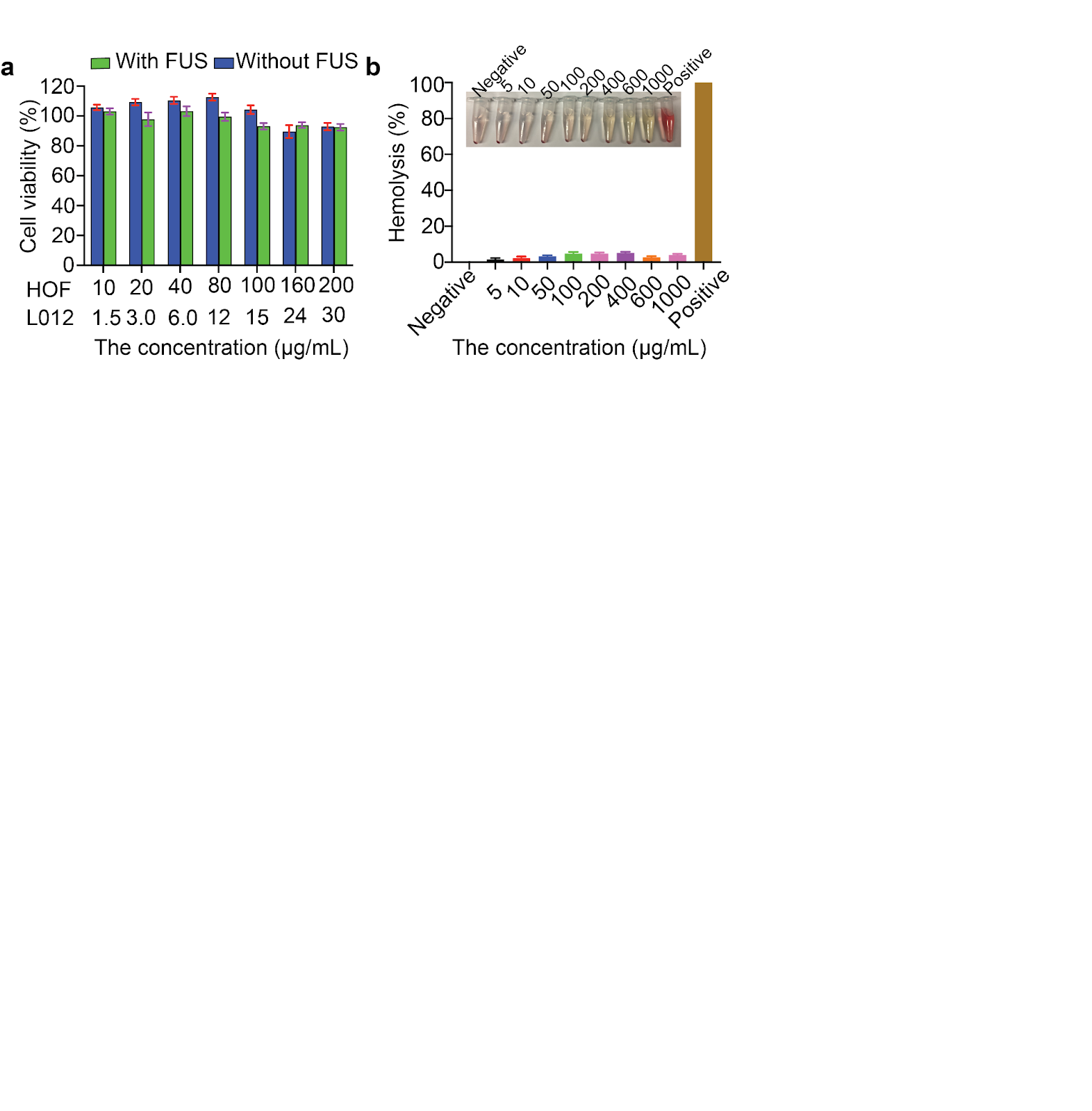

**Supplementary Fig. 10. The biosafety tests of HOF@L012 nanoparticles**. (a) The cell viability tests of HOF@L012 nanoparticles in HEK-293T cells with and without ultrasound stimulation. (b) The hemolysis tests of HOF@L012 nanoparticles.

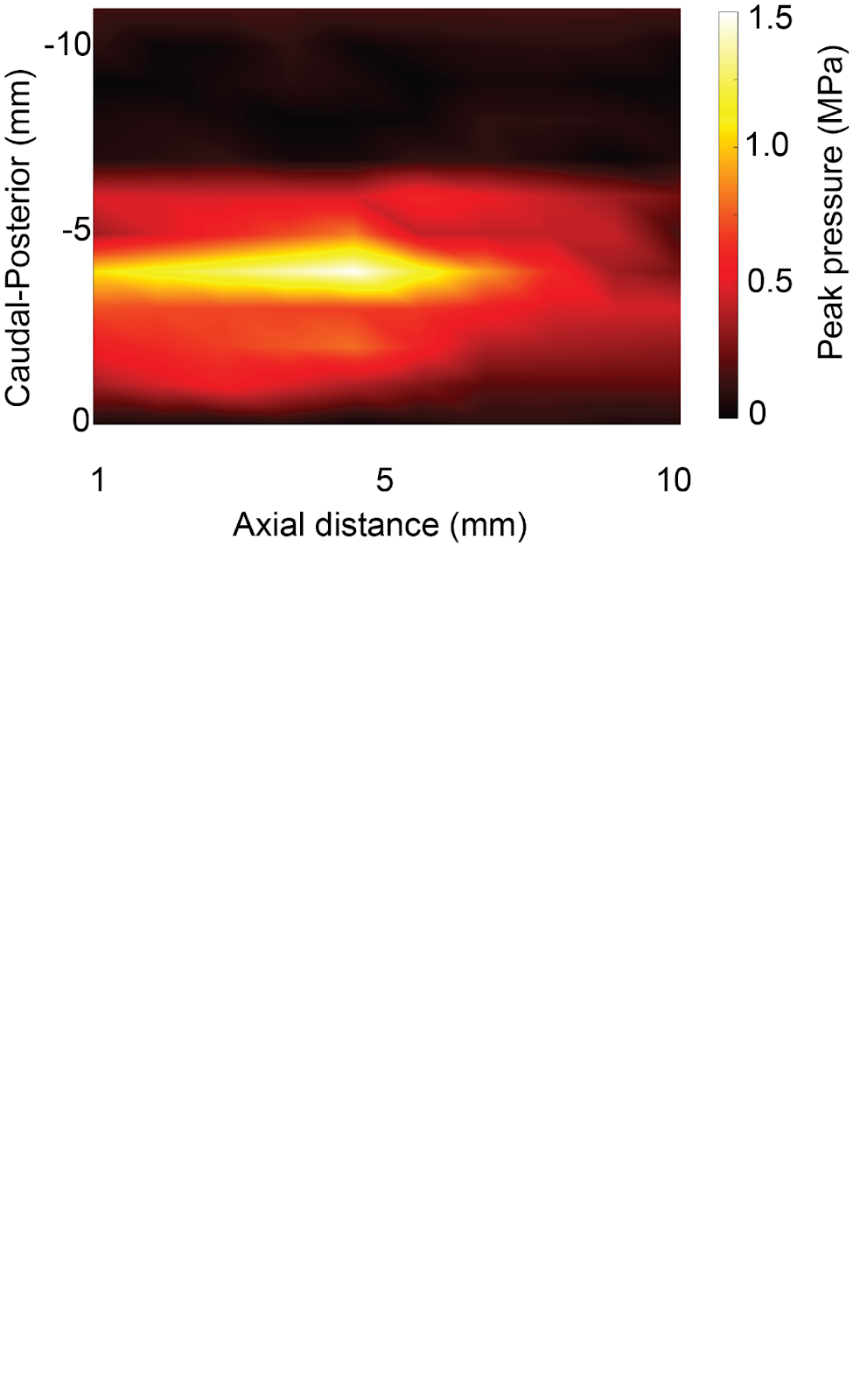

**Supplementary Fig. 11. The *in vivo* ultrasound power transfer in rat heads with FUS focus length of 5 mm.** The ultrasound power heatmap in the rat head shows that around 1.50 MPa was delivered to the rat GPe when 2.45 MPa primary ultrasound power was used.

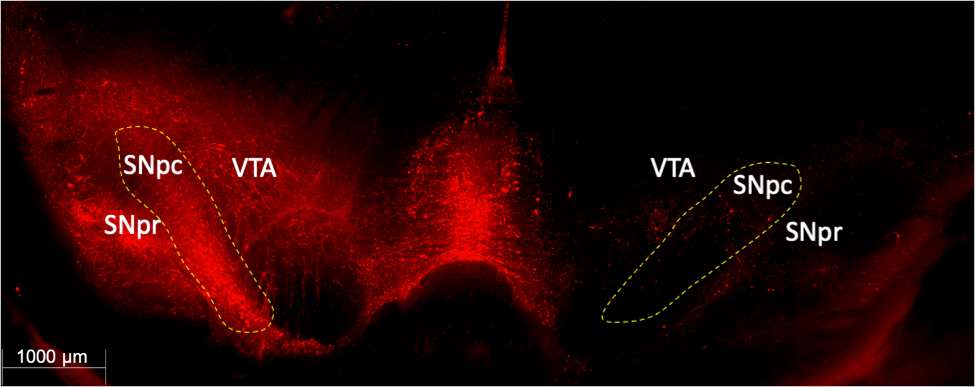

**Supplementary Fig. 12. Tyrosine hydroxylase (TH) immunofluorescence staining in hemiparkinsonian rat’s basal ganglia.** Unilateral 6-OHDA lesion resulted in dopamine neuron degeneration and death in the right substantia nigra pars compacta (SNpc) and surrounding regions with a significant decrease of TH+ dopamine neurons. *VTA (ventral tegmental area), SNpr (substantia nigra pars reticulata).*

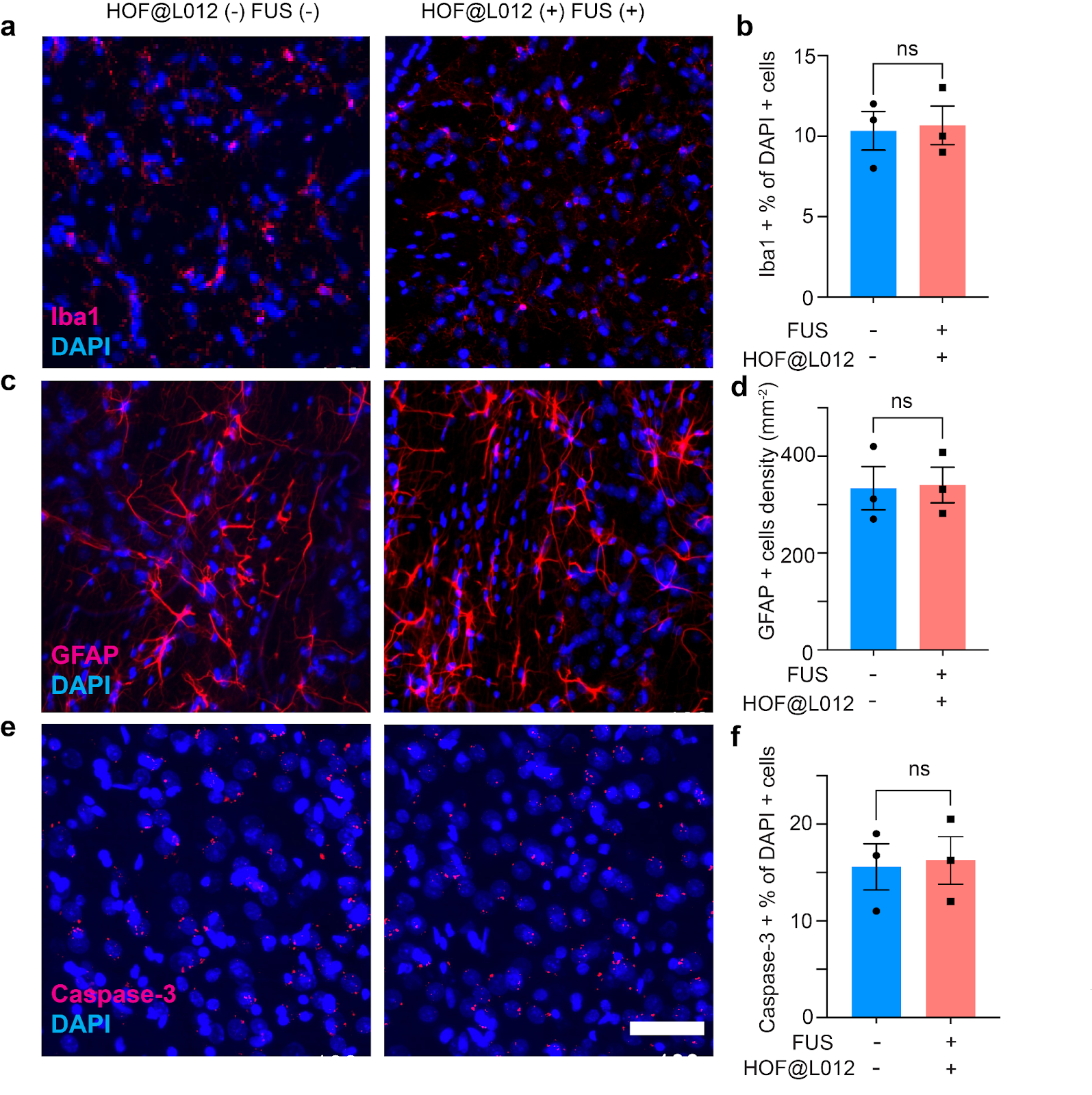

**Supplementary Fig. 13. *In vivo* biocompatibility evaluation of the sono-chemogenetics via determining microglia (Iba1) and astrocytes (GFAP) activation and neuronal apoptosis (Caspase-3).** Fluorescence images of Iba1 (a), GFAP (c), and Caspase-3 (e) of the brain slices in rat GPe brain region without stimulation and 7 days after sono-optogenetic stimulation. Scale bar: 50 μm. Blue signal is DAPI and the red signal is Iba1 (a), GFAP (c) and Caspase-3 (e). (b,d,f) Statistical analysis of the Iba1 (b), GFAP (d) and Caspase-3 (f) intensity. Mean ± SEM, n = 3 rats in each group. Two-way ANOVA and Tukey’s multiple comparison tests (P ≥ 0.05 (ns), * 0.01≤ P < 0.05, ** 0.001≤ P < 0.01, **** P< 0.0001).

**Tables**

**Supplementary Table 1.** Dynamic light scattering tests and drug loading content tests of various sono-HOF nanoparticles.

| entries | Nanoparticles | Size  (d.nm) | PDI | Zeta potential  (mv) | DLC  (wt%) |
| --- | --- | --- | --- | --- | --- |
| 1 | Blank HOF | 533.7±14.9 | 0.205 | -53.9±0.9 | N/A |
| 2 | HOF + 10% FBS | 538.3±11.6 | 0.452 | -22.2±1.5 | N/A |
| 3 | HOF@L012 | 525.7 ± 20.8 | 0.357 | -47.8±0.5 | 25.3±0.6 |
| Previous work^1^ | Lipo@IR780/L012 | 120.9 ± 0.5 | 0.189 | -25.8±0.7 | 4.3±0.4 |

**Supplementary Table 2.** Antibodies used in this work.

| **Primary antibodies** | **Secondary antibodies** |
| --- | --- |
| Rabbit anti-Iba1  (1:1000, 013-27691, Wako Chemicals) | Donkey anti-Rabbit, Alexa Fluor 594  (1:500, A32754, Invitrogen) |
| Rabbit anti-Cleaved Caspase-3  (1:1000, 9661, Cell Signaling Tec.) | Donkey anti-Rabbit, Alexa Fluor 594  (1:500, A32754, Invitrogen) |
| Rabbit anti-GFAP  (1:1000, 13-0300, Invitrogen) | Donkey anti-Rabbit, Alexa Fluor 594  (1:500, A32754, Invitrogen) |
| Mouse anti-tyrosine hydroxylase antibody (MA1-24654, Fisher Scientific, 1:1000) | Goat anti-mouse Alexa Fluor 488     (1:1000, ab150113, Abcam) |
| Rabbit anti-c-Fos antibody for mice  (1:500, ab222699, Abcam) | Goat anti-rabbit Alexa Fluor 594        (1:500, ab175652, Abcam) |
| Rabbit anti-c-Fos antibody for rats    (1:500, ab289723, Abcam) | Goat anti-rabbit Alexa Fluor 647         (1:500, A32733, Fisher Scientific) |
| Mouse anti-Parvalbumin antibody (P3088-100UL, Sigma-Aldrich, 1:1000) | Goat anti-mouse Alexa Fluor 594, (1:1000, A21125, Invitrogen) |
|  | Hoechst 33342  (1:5000, 17535, AAT Bioquest ) |

1.	Wang, W. *et al.* Ultrasound-Triggered In Situ Photon Emission for Noninvasive Optogenetics. *J. Am. Chem. Soc.* **145**, 1097–1107 (2023).
